# Lamin A/C Deficiency Drives Genomic Instability and Poor Survival in Small-Cell Lung Cancer through Increased R-loop Accumulation

**DOI:** 10.1101/2025.04.29.651052

**Authors:** Christopher W. Schultz, Sourav Saha, Anjali Dhall, Yang Zhang, Parth Desai, Lorinc S. Pongor, David A. Scheiblin, Valentin Magidson, Yilun Sun, Christophe Redon, Suresh Kumar, Manan Krishnamurthy, Henrique B. Dias, Vasilisa Aksenova, Elizabeth Giordano, Nobuyuki Takahashi, Michael Nirula, Mohit Arora, Chiori Tabe, Maria Thomas, Rajesh Kumar, Yasuhiro Arakawa, Ukhyun Jo, Beverly A. Teicher, Mirit I. Aladjem, Stephen Lockett, Mary Dasso, Yves Pommier, Ajit K. Sharma, Anish Thomas

**Author notes:** Corresponding Author: Anish Thomas. **Email:**. **Competing interests:** Authors declare that they have no competing interests. **Data and materials availability:** Upon publication all data will be uploaded to an appropriate platform.

## Abstract

*Lamin A/C* (*LMNA*), a key component of the nuclear envelope, is essential for maintaining nuclear integrity and genome organization [1]. While *LMNA* dysregulation has been implicated in genomic instability across cancer and aging, the underlying mechanisms remain poorly understood [2]. Here, we investigate *LMNA*’s role in small-cell lung cancer (SCLC), a highly aggressive malignancy characterized by extreme genomic instability [3, 4]. We demonstrate that *LMNA* depletion promotes R-loop accumulation, transcription-replication conflicts, replication stress, DNA breaks, and micronuclei formation. Mechanistically, *LMNA* loss disrupts nuclear pore complex distribution, reducing phenylalanine-glycine (FG)-nucleoporin incorporation and impairing RNA export efficiency. Furthermore, we show that *LMNA* expression is epigenetically repressed by *EZH2* during SCLC differentiation from neuroendocrine (NE) to non-NE states. Clinically, low *LMNA* levels correlate with significantly worse survival in SCLC patients. These findings uncover a novel role for *LMNA* in safeguarding genome integrity and shaping tumor heterogeneity, with broad implications for cancer and aging.

**Significance Statement:** Lamin A/C, a key structural component of the nuclear envelope, is frequently lost or mutated in cancer, laminopathies, and aging-related disorders. Lamin A/C loss is associated with genomic instability, but the underlying mechanisms remain incompletely understood. We demonstrate that LMNA loss drives genomic instability by promoting R-loop accumulation through disrupted nuclear pore dynamics and impaired RNA export. These findings reveal a previously unrecognized link between LMNA loss, nuclear envelope dysfunction, and genome instability. Targeting this pathway could help mitigate genomic instability in aging and laminopathies, while leveraging R-loop accumulation may enhance the efficacy of DNA-damaging therapies in cancer.

## Introduction

Lamins are a network of intermediate filaments localized at the inner nuclear membrane, comprising A-type lamins (lamin A/C) and B-type lamins (LMNB1, LMNB2) [1]. A-type lamins, which arise from alternative splicing of the *LMNA* gene, include two main isoforms, lamin A and lamin C. These isoforms are crucial for maintaining the structural integrity of the nucleus and play roles in chromatin organization, transcriptional regulation, nuclear pore complex (NPC) positioning, and cytoskeletal stability [1, 5]. Mutations or loss of A-type lamins, especially lamin A, have been linked to a range of diseases, including various cancers and laminopathies, such as cardiac and skeletal myopathies, lipodystrophies, and premature aging conditions including Hutchinson-Gilford progeria syndrome [6, 7].

A hallmark of both cancer and aging is genomic instability, which contributes to tumorigenesis and age-related diseases [8-12]. Despite this well-known association, the mechanisms by which lamin A/C loss contributes to genomic instability remain poorly understood [2]. One potential contributor is the accumulation of R-loops – three-stranded RNA/DNA hybrids that form during transcription [13, 14]. While R-loops are a normal part of cellular processes, their accumulation can lead to DNA damage, especially during DNA replication, where collisions between replication forks and R-loops can result in double-strand breaks (DSBs) [15]. Elevated R-loops have been observed in both aging and cancer [13, 14, 16, 17], and altered R-loop dynamics are implicated in LMNA-related diseases like lipodystrophy [18, 19]. Given the known relationship between R-loops and DNA damage [20], we hypothesized that R-loop accumulation may drive genomic instability in diseases involving lamin A/C loss. Small cell lung cancer (SCLC), which is characterized by generally low LMNA expression [21] and high genomic instability [3, 4], provides an ideal model for testing hypothesis.

### Lamin A/C suppresses R-loops and DNA damage

To investigate whether the genomic instability associated with *LMNA* loss is due to R-loops, we utilized the SCLC cell line DMS114 which expresses high levels of lamin A/C and, and as we have previously shown, exhibits low genomic instability [3]. Transient knockdown of *LMNA* (siRNA) resulted in more than a 2.2-fold increase in R-loops compared to control, as evaluated by slot-blot [15] (**Fig. 1A**). Similarly, *LMNA* knockout (KO) led to a 3.6-fold increase in R-loops (**Fig. 1A**).

**Figure 1:**
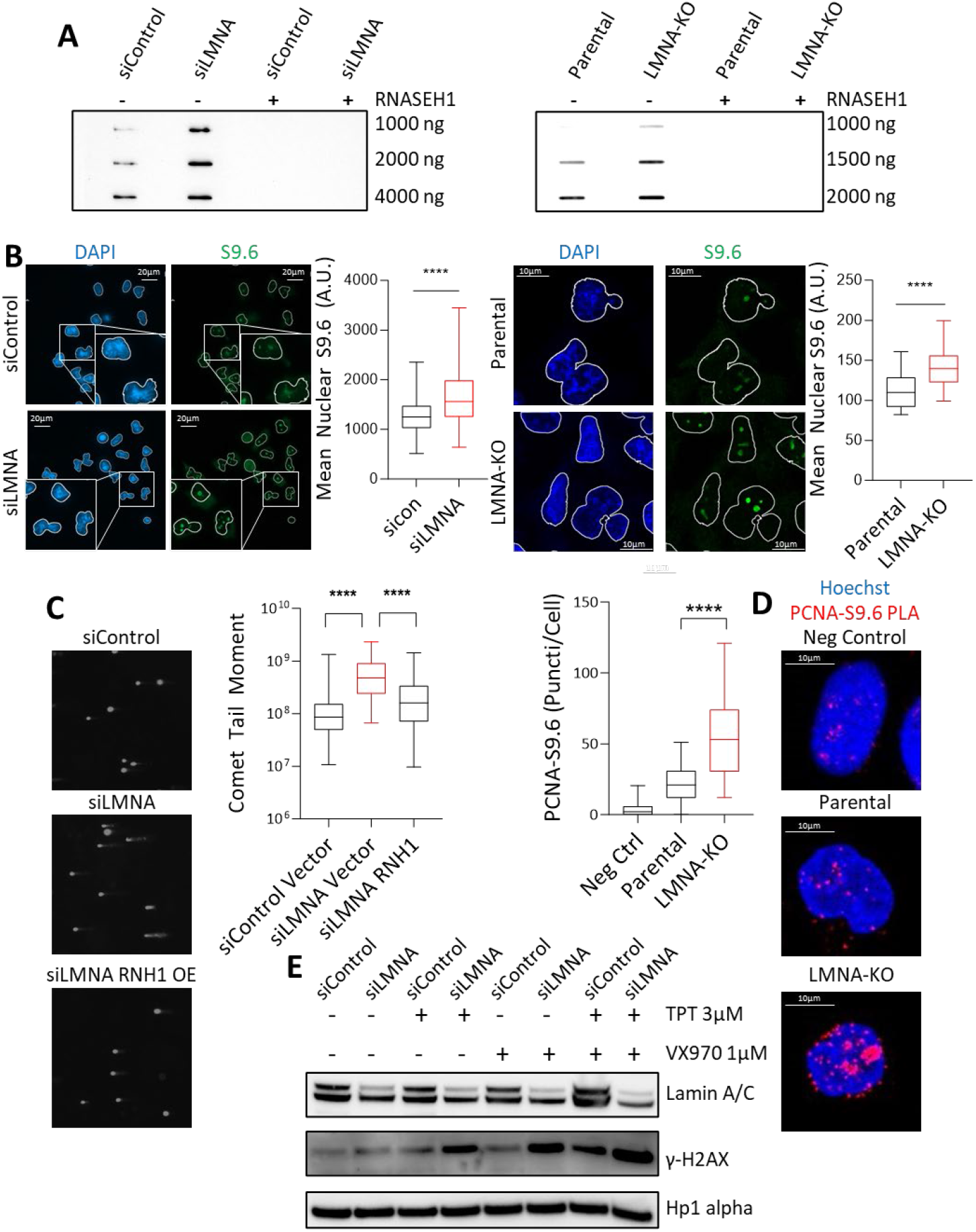
Lamin A/C suppresses R-loops and DNA damage: **A)** Slot blot of genomic DNA showing R-loops in siControl and siLMNA (left), parental and *LMNA*-KO (right) DMS114 cells, with and without RNaseH1. **B)** Representative immunofluorescence images of R-loops in siLMNA (left) and *LMNA*-KO (right) DMS114 cells. **C)** DNA damage detected by alkaline comet assay in siLMNA cells, with and without RNaseH1 in DMS114 cells. **D)** Proximity ligation assay assessing replication fork (PCNA) and R-loops (S9.6) in parental and *LMNA*-KO DMS114 cells. **E)** Immunoblotting of γH2AX to evaluate DNA damage after treatment with TOP1 inhibitor topotecan, ATR inhibitor berzosertib (VX-970), or co-treatment with both agents in siLMNA and siControl DMS114 cells.

To validate the observed increase in R-loops, we utilized immunofluorescence as a parallel approach [22]. Loss of lamin A/C using both siRNA and KO resulted in significant increases in R-loop signals (**Fig. 1B)**. Accumulation of R-loops following lamin A/C loss was observed throughout the nucleus, with predominant localization in the nucleoli. This was accompanied by an increase in nucleolar area, as indicated by nucleolin staining, consistent with *LMNA* loss-induced nucleolar expansion [23] (Fig. S1A). Overexpression of RNaseH1, a nuclease that specifically degrades RNA in RNA-DNA hybrids or overexpression of LMNA-GFP effectively suppressed R-loop formation in *LMNA*-KO cells (Fig. S1B-D). Consistent with these results, SCLC cell line NCI-H69, with lower basal levels of lamin A/C, exhibited more R-loops than DMS114; these effects were mitigated by LMNA-GFP overexpression (Fig. S1B, C).

R-loops are a known source of DNA damage and genomic instability [15]. We observed a significant increase in DNA breaks in siLMNA cells, as assessed by alkaline comet assay. These cells displayed longer comet tails with greater intensity after single-cell gel electrophoresis, indicating accrued DNA breaks. Importantly, this damage was linked to R-loop accumulation, as overexpression of RNaseH1, which resolves R-loops, reduced these features (**Fig. 1C**). Further, lamin A/C loss resulted in significantly elevated levels of DNA DSBs, as demonstrated by increased phosphorylated H2AX levels in siLMNA cells compared to control cells. DSB accumulation was also mitigated by RNaseH1 overexpression (Fig. S1E). Additionally, lamin A/C loss led to a marked increase in micronuclei formation (Fig. S1F), a hallmark of replication stress and genomic instability [24]. This effect too was reversed by RNaseH1 overexpression (Fig. S1G), further supporting the role of R-loops in driving genomic instability in *LMNA*-deficient cells. The increased replication stress with LMNA loss was corroborated by slower replication rates (Fig. S1H) and a decreased percentage of S-phase cells in *LMNA*-knockdown cells, an effect that was similarly rescued by RNaseH1 overexpression (Fig. S1I). These findings strongly suggest that R-loop formation constitutes a threat to genome stability in lamin A/C deficient cells.

Mechanistically, R-loop-mediated DNA damage can occur due to collisions between the transcription machinery and DNA replication forks [15]. To investigate whether lamin A/C loss leads to replication fork collisions via R-loop formation, we employed proximity ligation assay (PLA) [15] using antibodies specific for R-loops and proliferating cell nuclear antigen (PCNA), a key component of the replication fork. *LMNA*-KO cells showed a 2.7-fold increase in PLA signals compared to parental control cells, with a notable concentration of R-loop-PCNA foci in the nucleolar regions (**Fig. 1D**), suggesting that R-loop accumulation following lamin A/C loss disrupts normal replication processes, leading to fork stalling or collisions.

R-loop accumulation and the resultant genomic instability is counteracted by topoisomerase I (TOP1) which resolves DNA supercoiling and prevents R-loop formation [24]. Treatment with the TOP1 inhibitor topotecan significantly increased micronuclei formation in *LMNA*-KO cells compared to controls (Fig. S1F). Ataxia telangiectasia and Rad3-related (ATR) kinase reduces R-loop dependent DNA damage by suppressing transcription-replication collisions, promoting replication fork recovery, and enforcing G2/M checkpoint arrest [25]. A synthetically lethal combination of TOP1 and ATR inhibitors, currently in clinical development [26, 27], caused significantly more DNA DSBs in si*LMNA* cells than either agent alone (**Fig. 1E**). Collectively, these findings suggest that lamin A/C serves as a protective factor against R-loop accumulation, along with the ensuing DNA damage and genomic instability. Deregulation of lamin A/C expression may contribute to genomic instability.

### Lamin A/C loss induces R loops through disrupted RNA export

To investigate how lamin A/C loss mediates R-loop formation, we performed RNA-seq on parental and *LMNA*-KO cells. Compared to parental cells, *LMNA*-KO cells exhibited increased expression of genes associated with neuronal functions (e.g. *CDH7, BATF3*, and *MAP2*) (**Fig. 2A**) and decreased expression of genes linked to cardiac development (e.g. *SIPR1, PDE4D*) and cellular structural organization (e.g. *ACTA2, PDLIM1*) (**Fig. 2A**). These findings were further supported by single-sample Gene Set Enrichment Analysis (ssGSEA) and GSEA, which revealed enrichment of pathways related to neuronal differentiation (e.g. noradrenergic neuron development) and depletion of pathways related to cardiac development (e.g. heart formation), heterochromatin organization (e.g. chromosome attachment to the nuclear envelope), cellular structural organization (e.g. regulation of microtubule nucleation), and additionally nuclear pore dynamics (e.g. nuclear pore), and RNA transport (e.g. mRNA transport) in *LMNA*-KO cells (**Fig. 2B, C**).

**Figure 2:**
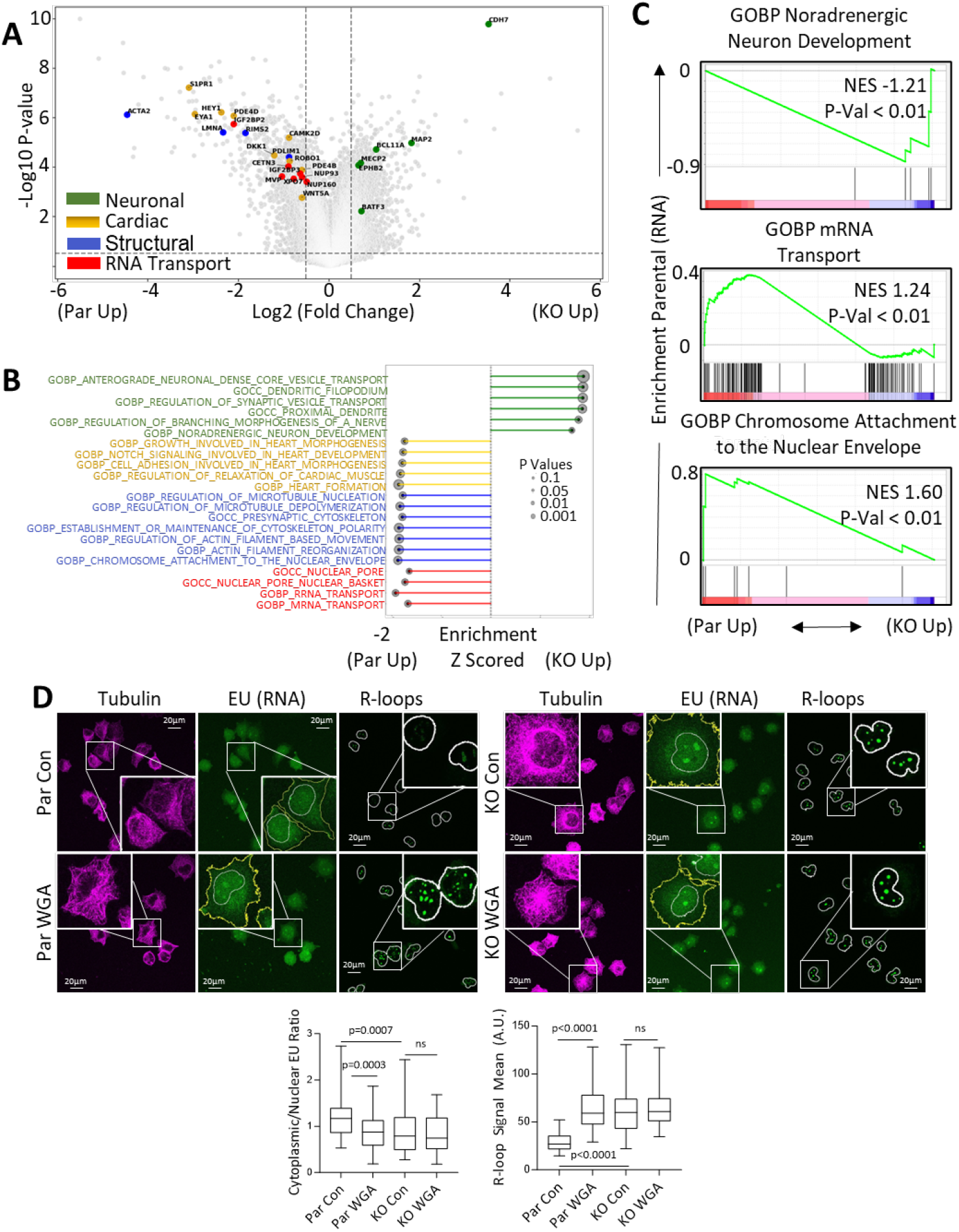
Lamin A/C loss induces R loops through disrupted RNA export: **A)** Volcano plot showing differentially expressed genes between parental and *LMNA*-KO DMS114 cells. **B, C)** Pathway enrichment in parental vs. *LMNA*-KO DMS114 cells as assessed by ssGSEA (B) and GSEA (C). **D**) RNA export and R-loop formation assessed following WGA treatment in parental and *LMNA*-KO cells. RNA export was measured by EU incorporation, and R-loops detected by immunofluorescence, representative images are shown with nuclei outlined in white while the cytoplasm is outlined in yellow.

To assess if these changes in gene expression were due to altered gene accessibility, we performed ATAC-seq. *LMNA*-KO cells exhibited greater accessibility near transcription start sites (TSS), leading to an overall increased accessibility of promoter regions, compared to control cells (Fig. S2A, B). ATAC-seq confirmed increased accessibility of genes related to neuronal differentiation, while genes related to cellular structural organization, cardiac development and function, as well as RNA export showed reduced accessibility (Fig. S2C-E). These findings support previous reports linking LMNA loss to overexpression of neuronal genes and altered heterochromatin organization [28], as well as the low LMNA expression in neuronal lineages (Fig. S2F) [29]. Lamin A/C plays a key role in nuclear envelope integrity through its interaction with the LINC complex, which maintains nuclear and cytoskeletal stability. Mutations in *LMNA* cause mechanically unstable nuclei, contributing to myopathies [30, 31].

Given the depletion of RNA export in *LMNA*-KO cells, and the known link between RNA export and R-loop formation [22, 32-35], we evaluated RNA export using 5-ethynyl uridine (EU) labeling to track nascent RNA distribution [36] between the nucleus and cytoplasm. Loss of lamin A/C resulted in significant nuclear retention of nascent transcripts (Fig. S2G). To test whether RNA export inhibition could induce R-loops, we treated parental and *LMNA*-KO cells with wheat-germ agglutinin (WGA), a nucleoporin-binding agent that inhibits RNA export [37, 38]. In parental cells, WGA treatment inhibited RNA export and induced R-loops. However, in *LMNA*-KO cells, WGA neither further inhibited RNA export nor increased R-loop formation (**Fig. 2D**). This may reflect a ceiling effect, where RNA export is already maximally impaired due to disrupted nuclear pore function in LMNA-KO cells [38]. Alternatively, both WGA and LMNA depletion may converge on a shared pathway regulating nuclear pore dynamics, preventing an additive effect on RNA export and R-loop formation.

To further confirm the link between RNA export inhibition and R-loop formation, we transiently silenced the highly conserved RNA export factors *TPR* and *NXT1* [22, 39] in parental DMS114 cells. Knockdown of NXT1, TPR, and LMNA all significantly reduced RNA export, and consistent with the role of RNA export in regulating R-loop formation [22, 33-35], this led to a marked increased R-loop formation (Fig. S2H). Together, these results suggest that lamin A/C facilitates the nuclear export of RNA transcripts, thereby maintaining R-loop homeostasis.

### Lamin A/C loss disrupts RNA export and causes R loops by altering the distribution of nucleoporins

We performed DRIP-seq (DNA:RNA immunoprecipitation followed by sequencing) to map R-loop formation genome-wide in both parental and *LMNA*-KO cells. Consistent with our earlier findings (Fig. 1), we found a significant increase in R-loop peaks in *LMNA*-KO compared to parental cells (**Fig. 3A**, S3A). Interestingly, these R-loops were predominantly enriched in genic regions (**Fig. 3B**, while intergenic R-loop formation remained unchanged (Fig. S3B). This contrasts with another R-loop suppressor TOP1, which does not alter the genomic distribution of R-loop peaks [40], further supporting the role of altered RNA export in R-loop formation upon LMNA loss.

**Figure 3.**
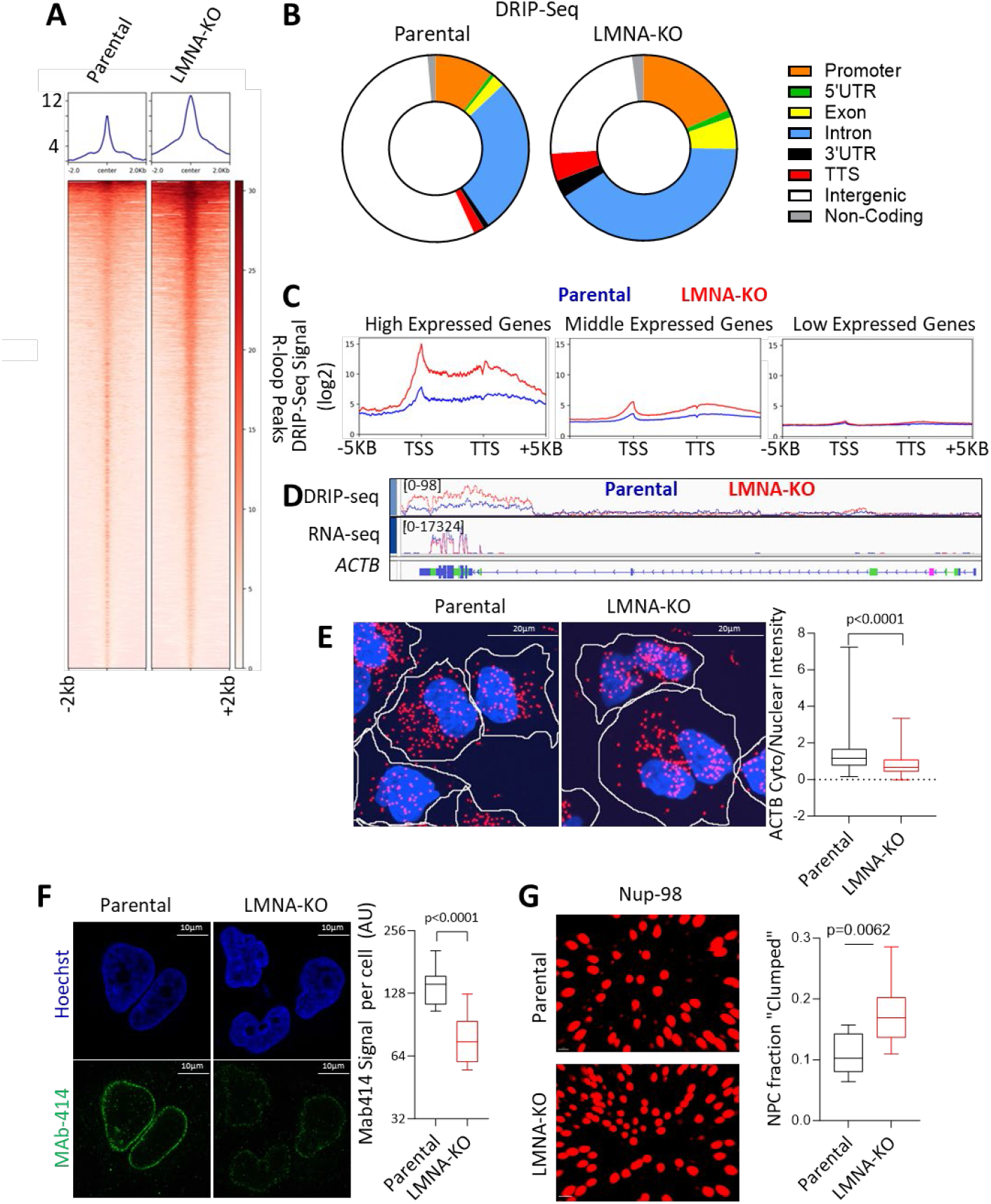
Lamin A/C loss disrupts RNA export and causes R loops by altering the distribution of nucleoporins. **A)** DRIP-seq signal intensity in parental vs *LMNA*-KO DMS114 cells. **B)** R-loop peaks analyzed across parental and *LMNA*-KO DMS114 cells, with peaks assigned to gene categories based on the most proximal transcription start site. **C)** Average DRIP-seq signal across gene length for high (>6 log2 TMM FPKM, 447 genes), medium (1-5 log2 TMM FPKM, 10,045 genes), and low (<1 log2 TMM FPKM, 27,290 genes) expression categories, as determined by RNA-seq data. **D)** IGV tracks, alignment of DRIP-seq and RNA-seq signals for the highly transcribed gene *ACTB*. **E)** FISH analysis of the cytoplasmic versus nuclear distribution of *ACTB* RNA in parental and *LMNA*-KO cells. Cytoplasmic borders are outlined in white. **F)** FG nucleoporins assessed by immunofluorescence using MAb414 staining in parental and *LMNA*-KO cells. **G)** NUP98 staining and expansion microscopy to assess for NPC distribution. NPC’s which have an outer rim diameter of ∼120 nm [42] were considered “clumped” if they were within 180 nm.

In *LMNA*-KO DMS 114 cells, R-loop peak intensity was markedly higher, with an 11.6-fold increase, in highly expressed genes (**Fig. 3B**, S3C). Medium-expressed genes showed a 2.03-fold increase, while low-expressing genes had a modest 1.1-fold increase. R-loop enrichment was particularly pronounced at transcription start sites (TSS) and transcription termination sites (TTS) (**Fig. 3C**), known hotspots for RNA Pol-II pausing and R-loop formation [41]. In low-expressing genes, the signal at TSS and TTS was stronger relative to the central gene body (Fig. S3D), while highly expressed genes maintained sustained R-loop signals between TSS and TTS peaks. RNA-fluorescence in situ hybridization (FISH) of the highly transcribed gene *ACTB* confirmed reduced RNA export in *LMNA*-KO cells, as evidenced by a decreased cytoplasmic/nuclear ratio of *ACTB* RNA (**Fig. 3D, E**). These findings align with previous studies linking RNA export inhibition to increased R-loop formation [22, 33-35], particularly in highly transcribed genes [33, 40].

RNA export relies on NPCs, composed of nucleoporins that form channels between the nucleus and cytoplasm [42]. Loss of lamin A/C disrupts the localization of FG nucleoporins [43], specialized nucleoporins containing phenylalanine-glycine (FG) repeat domains that are essential for NPC interaction with RNA [42]. Given the central role of lamin A/C in maintaining nuclear architecture [1], we hypothesized that reduced RNA export in *LMNA*-KO cells might be due to altered NPC dynamics. Immunostaining with the MAb414 antibody [43], which primarily targets FG nucleoporins, showed diminished staining at the nuclear periphery in *LMNA*-KO cells (**Fig. 3F**), suggesting either a decrease in NPC number or reduced incorporation of FG nucleoporins into individual NPCs. However, high-resolution SIM microscopy indicated no significant difference in the number of NPCs, supporting that the reduced MAb414 signal is due to reduced incorporation of one or more FG-nucleoporins into NPCs (Fig. S3E, F).

Immunoblot analysis of FG nucleoporins using MAb414 demonstrated lower expression of nucleoporins Nup153 and Nup62 in *LMNA*-KO cells, while Nup98 levels remained unchanged (Fig. S3G). Further validation with a Nup153 antibody confirmed a near-complete loss of Nup153 protein (Fig. S3H). The reduction of individual FG nucleoporins is critical as it can impair RNA export [44]. Importantly, lamin A/C directly interacts with Nup153 to facilitate its localization [45] and loss/inhibition of Nup153 alone can result in decreased nucleoporin incorporation into NPCs, NPC clustering, and reduced mRNA export [46, 47].

To examine the distribution of NPCs in the context of lamin A/C deficiency, we utilized expansion microscopy to preserve structural relationships while staining for Nup98, which remained stable in *LMNA*-KO cells. Our analysis revealed a significant increase in the proportion of highly proximal NPCs, indicating NPC clustering (**Fig. 3G;** Fig. S3I, J). We assessed *LMNA* expression in cancer cell lines and observed a strong negative correlation between expression of *LMNA* and most nucleoporins (Fig. S3K). This finding potentially indicates a compensatory up-regulation of nucleoporins in low *LMNA* cells in response to reduced FG nucleoporin incorporation and NPC clumping. In summary, loss of lamin A/C disrupts RNA export efficiency by altering NPC distribution and reducing the incorporation of select FG nucleoporins, leading to a substantial increase in intragenic R-loops, particularly in highly transcribed genes.

### EZH2-mediated regulation of *LMNA* and SCLC differentiation

A-type lamins have important roles in differentiation and are generally absent or lowly expressed in undifferentiated cells and more highly expressed in differentiated cells [48, 49]. While SCLC tumors are predominantly neuroendocrine (NE), they also contain non-neuroendocrine (non-NE) cells, as shown in genetically engineered mouse models, patient-derived xenografts, and human tumors [50]. NE cells can differentiate into non-NE cells, losing NE transcription factor expression and gaining non-NE markers in a process driven by MYC and NOTCH [51]. NE-to-non-NE transition is associated with distinct structural changes, such as a shift from suspension to adherent growth in cultured cells [50, 52, 53]. Given these changes, we hypothesized that *LMNA* expression might be dynamically modulated during SCLC differentiation from an NE to non-NE state.

Time-series transcriptome analysis of a MYC and NOTCH-driven genetically engineered mouse model of SCLC [51] revealed an increase in *LMNA* expression coinciding with SCLC differentiation to a non-NE state (Fig. S4A). This was further supported by a significant negative correlation between NE gene expression and *LMNA* levels in SCLC cell lines and human tumors (Fig. S4B). To validate this relationship, we assessed INSM1 expression and suspension culture growth as markers of NE differentiation [50, 53]. In a panel of human SCLC cell lines with varying NE differentiation levels, INSM1 was highly expressed in NE cells and nearly absent in non-NE cells. Correspondingly, lamin A/C expression and adherent growth were significantly increased in non-NE cells (**Fig. 4A**, S4C).

**Figure 4:**
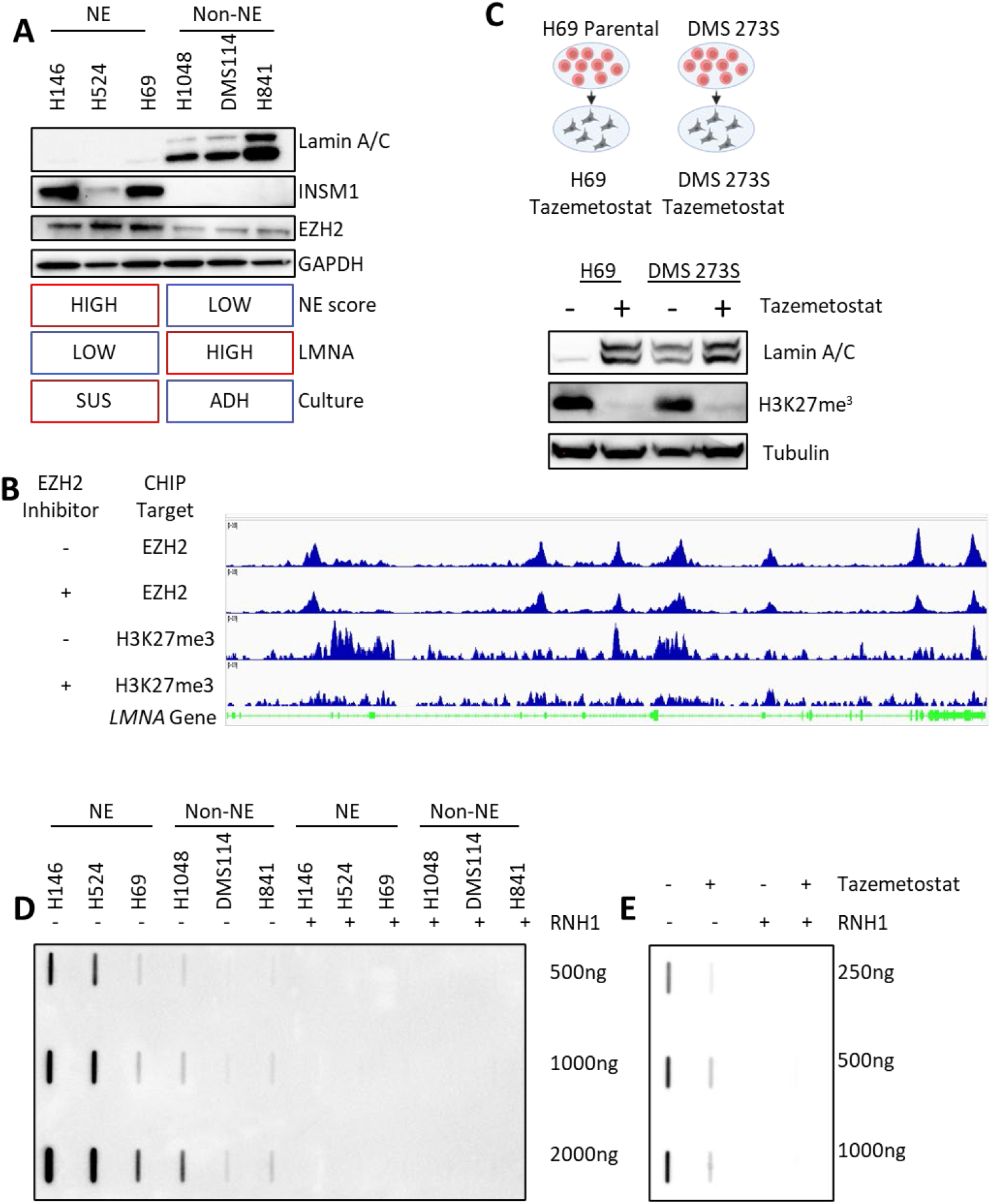
EZH2-mediated regulation of LMNA and SCLC differentiation: **A)** Western blot analysis of INSM1 (NE marker), lamin A/C, and EZH2 in NE and non-NE SCLC cell lines. **B)** Integrated Genome Viewer tracks showing EZH2 and H3K27me3 chromatin immunoprecipitation (ChIP) data from an SCLC tumor model treated with vehicle or the EZH2 inhibitor EPZ011989. **C)** Lamin A/C and H3K27me3 levels in DMS273S and NCI-H69 cells after treatment with the EZH2 inhibitor tazemetostat. **D)** R-loop detection by slot blot in a panel of SCLC cell lines. **E)** R-loop detection by slot blot analysis in NCI-H69 cells treated with tazemetostat.

To further explore this dynamic, we used two isogenic models. In the NE cell line NCI-H69, doxorubicin treatment induces differentiation into non-NE H69AR cells [53]. H69AR cells exhibited the expected transition to adherent growth, loss of INSM1 expression, and a significant increase in lamin A/C expression (Fig. S4D). Similarly, in an isogenic model of DMS273, adherent non-NE (DMS273A) and suspension NE (DMS273S) subpopulations [54], DMS273A cells showed reduced INSM1 expression and ∼18-fold higher lamin A/C levels compared to DMS273S cells (Fig. S4E). These findings suggest that lamin A/C expression is upregulated during non-NE differentiation and may act downstream of factors such as MYC, NOTCH, or the NOTCH effector REST. Supporting this, LMNA-GFP overexpression in NCI-H69 cells partially reduced INSM1 expression but did not alter adherence, indicating that lamin A/C might contribute to, but is not solely responsible for, the transition (Fig. S4F). These findings align with those in neuroblastoma, which shares similar molecular characteristics with SCLC, where lamin A/C is crucial in the transition from NE to non-NE differentiation [48].

Unlike laminopathies, where LMNA mutations or copy number losses are common, such alterations are rare in SCLC (Table S1). Instead, we observed significant hypermethylation of the LMNA promoter in SCLC tumors compared to normal tissues, corresponding with reduced LMNA expression [55]. These findings implicate epigenetic regulation in the suppression of LMNA. Given the role of EZH2, a histone methyltransferase within the polycomb repressive complex 2, in driving methylation and NE differentiation in SCLC [55, 56], we hypothesized that EZH2 modulates LMNA expression. Analysis of SCLC xenograft data [56] confirmed EZH2 binding to the LMNA gene, and treatment with an EZH2 inhibitor in vivo reduced *LMNA* methylation (**Fig. 4B**). Consistently, in our cell line panel and isogenic models, EZH2 expression was elevated in NE cells and decreased upon non-NE differentiation (Fig. 4A, S4D, E).

To test whether pharmacological EZH2 inhibition could restore lamin A/C expression, we treated NCI-H69 and DMS273S cells with tazemetostat, an EZH2 inhibitor. Treatment reduced H3K27me3 levels, promoted an adherent phenotype (Fig. S4G), and induced up to a 30-fold increase in lamin A/C expression (**Fig. 4C**). Analysis across the Cancer Cell Line Encyclopedia (CCLE) confirmed a negative correlation between *LMNA* expression, NE differentiation, *LMNA* methylation, and *EZH2* expression, with SCLC cells uniquely characterized by low *LMNA* expression, high NE differentiation, and elevated *EZH2* expression and *LMNA* promoter methylation compared to other solid tumors (Fig. S4H, I).

Finally, we investigated whether the association between low *LMNA* expression and increased R-loops extended to NE differentiation models. Slot blot analysis confirmed that higher *LMNA* expression correlated with reduced R-loop formation across SCLC cell lines (**Fig. 4D**, Fig. S4J). In the DMS273 isogenic model, DMS273A cells displayed approximately four-fold fewer R-loops compared to DMS273S cells (Fig. S4K). Tazemetostat-treated NCI-H69 cells exhibited a five-fold reduction in R-loop formation alongside increased lamin A/C expression (**Fig. 4E**). Collectively, these data support a paradigm where NE differentiation drives increased R-loops in SCLC via EZH2 mediated epigenetic silencing of *LMNA*.

### LMNA loss is associated with NE differentiation, R-loop formation, and poor prognosis in SCLC patients

Human SCLC exhibits substantial heterogeneity, including both NE and non-NE cell populations [57]. To investigate whether the relationship between NE differentiation, lamin A/C expression and R-loops holds true in human SCLC, we performed multiplex immunofluorescence on tissue sections obtained from a rapid autopsy, staining for NE lineage-defining transcription factor ASCL1, lamin A/C, and R-loops (**Fig. 5A**, S5A). The analysis revealed that ASCL1 and lamin A/C expression were mutually exclusive, with ASCL1 present in 22.8% and lamin A/C in 17.2% of cells, with only 0.9% of cells exhibiting co-expression (**Fig. 5B**). This finding aligns with our earlier data showing that increased lamin A/C expression correlates with non-NE differentiation (Fig. 4). R-loops were most prevalent in ASCL1-positive cells, with a strong correlation between NE differentiation (ASCL1/lamin A/C ratio) and R-loop levels (P = 0.0207, R^2^ = 0.9577). This correlation was further confirmed in a tissue microarray panel of SCLC tumor samples (**Fig. 5C**, S5B-D).

**Figure 5:**
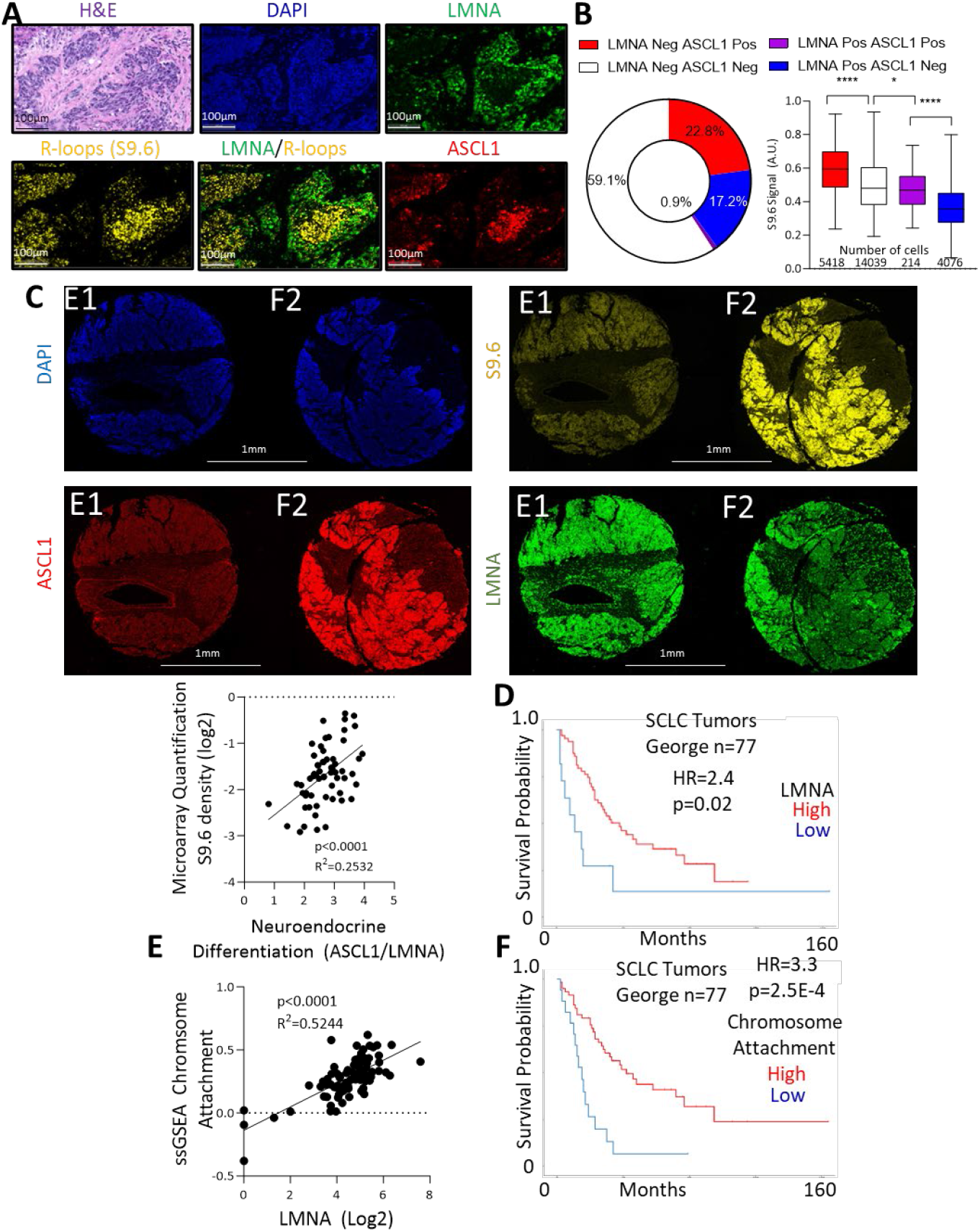
Lamin A/C loss is associated with NE differentiation, R-loop formation, and poor prognosis in SCLC patients: **A)** Rapid autopsy sample RA-17 stained for DAPI, ASCL1, LMNA, and R-loops, with accompanying H&E staining. **B)** Quantification of R-loop signals per cell in RA-17, grouped by ASCL1 (+/-) and LMNA (+/-) staining status. **C)** SCLC tissue microarray stained for DAPI, ASCL1, LMNA, and R-loops (S9.6), along with H&E staining. Two representative cores are shown (top), with analysis of ASCL1/LMNA signals and R-loop levels (bottom). **D)** Kaplan-Meier plot of overall survival stratified by LMNA RNA expression in SCLC patients [59]. **E)** Correlation between LMNA RNA expression and the “Chromosome Attachment to the Nuclear Envelope” gene set in SCLC tumors [59]. **F)** Kaplan-Meier plot of overall survival stratified by expression of the “Chromosome Attachment to the Nuclear Envelope” gene set in a human SCLC dataset [59].

Interestingly, lamin A/C-negative tumor regions often exhibited indistinct nucleoli (Fig. S5E), a known histological hallmark of SCLC [50]. Re-evaluating the expansion microscopy data, we noted a substantial reduction of the clear DNA border at the periphery of nucleoli in *LMNA* KO compared with parental cells (Fig. S5F). To validate this observation, we performed additional H&E staining, confirming that lamin A/C loss contributes to the reduced visibility of nucleoli (Fig. S5G), which aligns with previous studies showing diminished nucleolar definition with LMNA KO [58]. These findings reinforce the role of low LMNA expression in driving this characteristic histologic feature of SCLC.

To test whether lamin A/C-loss induced R-loops might contribute to poor prognosis, we first assessed whether *LMNA* RNA expression could predict protein levels in 63 SCLC cell lines, finding a positive correlation between RNA and protein expression (Fig. S5H,I, Table S2). Consistent with our hypothesis, low *LMNA* RNA expression was associated with poorer overall survival in a cohort of primary SCLC patients (HR=2.4, p=0.02, median 38 vs. 11 months) (**Fig. 5D**) [59]. Additionally, a gene set related to lamin A/C function, including genes involved in chromosome attachment to the nuclear envelope (*LMNA, SUN1, IFFO1, SPDYA, MAJIN, TERB1, TERB2*), showed significant correlation with *LMNA* expression and was more predictive of poor overall survival (HR=3.3, p=2.53E-4, median 42 vs. 15 months) (**Fig. 5E,F**).

In line with the above observations, and further supporting the role of R-loops as drivers of poor prognosis, reduced expression of multiple well-established R-loop suppressor genes, including *RNASEH1, BRD4, SETX*, and *DHX9*, were associated with reduced survival, either individually or together with *LMNA* (HR=4.6, p=1.07E-5, median 42 vs. 13 months) (Fig. S5J,K) [59, 60]. Notably, our findings contrast from previous studies which have suggested that low expression of *DHX9* is associated with better prognosis in lung cancer [61], likely due to differences in lung cancer subtypes. Together, our findings link SCLC heterogeneity and lamin A/C loss to increased R-loops and poor prognosis in SCLC patients and suggest that reduced lamin A/C contributes to the inconspicuous or absent nucleoli, a histological hallmark of SCLC.

To explore the broader applicability of these findings, we analyzed data from the PARCB model, in which prostate cells are driven towards a small cell lineage through *RB1* and *TP53* inactivation, coupled with overexpression of *c*-*MYC, BCL2*, and myristoylated *AKT1* [62]. This model recapitulates small cell neuroendocrine prostate cancer (SCNPC), a rare initial presentation but a significant subset of resistant prostate adenocarcinomas (PRAD) [61]. As the model progressed, we observed NE differentiation increases accompanied by elevated EZH2 expression and reduced LMNA expression (Fig. S5L). Consistent with our findings in SCLC, analysis of TCGA PRAD tumors revealed that low LMNA and low RNASEH1 expression were both significantly associated with poor overall survival (Fig. S5M). These results suggest that the EZH2-driven suppression of LMNA and its association with R-loop accumulation and decreased survival may extend to other small cell cancer types.

### Lamin A/C deficiency promotes genomic instability in aging

We have shown that lamin A/C suppresses R loop formation and DNA damage in SCLC. Given that reduced LMNA is a common feature of cancer and aging cells [8, 63], we explored whether the phenotypes observed with LMNA loss in SCLC were also present in aged cells with diminished lamin A/C levels.

To investigate this, we analyzed a pair of fibroblast cell lines from the same donor collected at ages 48 and 63. We first examined H3K9me3, a heterochromatin marker known to decline with age [63, 64]. As expected, aged cells exhibited a substantial reduction in H3K9me3 levels (**Fig. 6A**) [63]. Next, to determine whether lamin A/C loss directly contributed to the reduction in H3K9me3, we examined H3K9me3 levels in our isogenic DMS114 SCLC model. Lamin A/C loss resulted in a moderate reduction in H3K9me3 (**Fig. 6A**, Fig.S5N), consistent with the role of lamin A/C in maintaining heterochromatin [8]. However, additional factors likely contribute to the more pronounced H3K9me3 reduction in aging cells [63].

**Figure 6:**
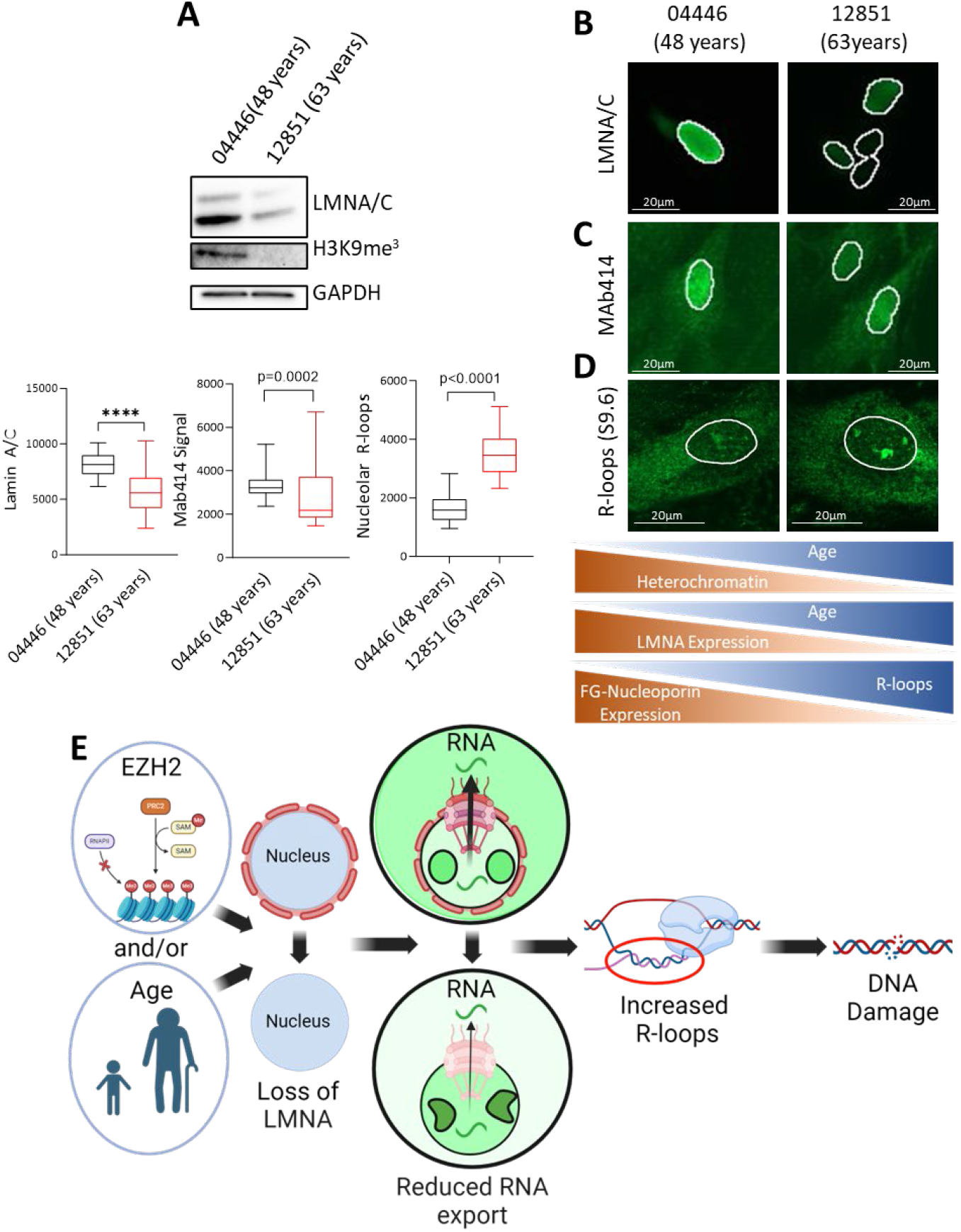
Lamin A/C deficiency promotes genomic instability in aging. **A)** H3K9me^3^ and lamin A/C assessed by immunoblot in patient-matched fibroblasts at 48-years and 63-years. **B)** Immunofluorescence for LMNA in the paired fibroblasts. **C, D)** Immunofluorescence for MAb414 (D) and R-loops/S9.6 (E) in younger (04446 48) and older (12851 63) fibroblasts. **E)** LMNA loss can occur due to various factors including EZH2 suppression and age. This results in reduced FG-nucleoporin incorporation in nuclear pores, leading to altered nuclear pore localization and decreased RNA export. These changes drive increased R-loop formation and subsequent DNA damage.

After confirming this hallmark of cellular aging, we assessed lamin A/C expression in older compared with younger fibroblasts. Immunostaining revealed a significant reduction in nuclear lamin A/C in aged cells, with a 4.1-fold decrease in total lamin A/C protein expression (**Fig. 6A, B**). We then examined whether this reduction in lamin A/C coincided with decreased FG-nucleoporin incorporation, as observed in our SCLC models. Older fibroblasts showed a significant reduction in nuclear MAb414 staining (**Fig. 6C**), consistent with previous findings of lower nucleoporin levels in aging cells [63] and our earlier results (Fig. 3). This reduction in lamin A/C and NPC integrity was associated with a pronounced increase in R-loop accumulation (**Fig. 6D**), consistent with prior reports of R-loop accumulation in highly transcribed genes with age [13], and our observations in lamin A/C-depleted SCLC.

Together our data indicate a central role for lamin A/C in safe-guarding cells from aberrant R-loop accumulation and genotoxic stress in multiple disease states. Loss of lamin A/C – whether through aging, or EZH2 mediated suppression as in the case of SCLC – leads to reduced FG-nucleoporin incorporation into NPCs, and disrupts RNA export, resulting in increased R-loops and genotoxic stress (**Fig. 6E**).

## Discussion

The importance of *LMNA*, which codes for lamin A and lamin C proteins, is underscored by its frequent alterations in cancer and aging-related disorders [6]. Lamin A/C provides structural support to the nucleus, organizes chromatin, and regulates gene expression, playing a crucial role in maintaining genomic stability. However, the specific mechanisms by which LMNA loss contributes to genomic instability, a hallmark of both cancer and aging, remain inadequately understood [2].

We find that: (i) Lamin A/C is essential for protecting cells from accumulation of R-loops, three-stranded nucleic acid structures that induce DNA damage and promote genomic instability. (ii) In SCLC, lamin A/C is silenced by the polycomb repressive complex 2 through EZH2, which trimethylates histone H3K27. Pharmacological inhibition of EZH2 reactivates LMNA expression, reducing R-loop accumulation and genomic instability while altering tumor cell fate. (iii) Loss of lamin A/C leads to increased R-loop formation, heightened genomic instability, and poor prognosis in SCLC patients. Reactivating LMNA offers a potential therapeutic strategy to address genomic instability in SCLC and potentially other LMNA-deficient conditions.

Previous studies focused on how lamin deficiencies disrupt conventional DNA replication and repair pathways [2]. Lamin A/C deficiency or mutations impair DNA repair by delaying the recruitment or causing the degradation of critical repair proteins such as FANCD2 [65] and 53BP1 [66, 67]. Moreover, disruptions in the nuclear lamina alter the spatial and temporal organization of DNA replication by mislocalizing key replication factors, including PCNA and RFC, which become trapped within aberrant lamin aggregates [68]. Our findings extend this body of knowledge by introducing R-loops as a new and important mechanism of DNA damage linked to *LMNA* loss.

Mechanistically, our findings reveal that *LMNA* loss disrupts the distribution and incorporation of key FG nucleoporins within NPCs, leading to structural alterations, including NPC clumping, which impair RNA transport. The mislocalization of FG nucleoporins likely promotes their degradation, a hypothesis that warrants further investigation [43, 65]. The disruption in RNA export is particularly deleterious for highly expressed genes, where defective RNA export exacerbates R-loop formation at transcriptional hotspots, such as transcription start and termination sites. These results align with the established role of *LMNA* in maintaining NPC integrity [43] and extend recent studies in a fly model of progeroid syndrome caused by A-type lamin mutations, where abnormal lamina organization was shown to impair the export of mitochondrial RNA, contributing to premature aging [32].

While our findings strongly support RNA export inhibition as the primary driver of R-loop induction associated with *LMNA* loss, they do not exclude other plausible mechanisms. *LMNA* deficiency could increase the accessibility of R-loop hotspots at TSS regions [8, 41]. However, this mechanism would likely lead to R-loop formation primarily at TSS regions of lowly transcribed genes, which is inconsistent with our data. Another possibility is that *LMNA* loss may indirectly downregulate *BRCA1*, a key regulator of R-loops [69, 70]. Yet, we observed no downregulation of *BRCA1* in *LMNA*-KO cells, and SCLC typically does not exhibit low *BRCA1* levels despite low *LMNA* expression.

Lamin A/C is broadly expressed in most somatic cells but notably absent or reduced in hematopoietic cells, neurons, undifferentiated epithelial and mesenchymal cells, and various cancers [7, 29, 48, 49]. In SCLC, LMNA expression inversely correlates with NE differentiation – higher NE marker levels coincide with lower LMNA expression. Increased LMNA levels in non-NE differentiation may drive the shift from suspension to adherent growth in cultured cells, as LMNA is critical for this process [71]. Our findings suggest that *LMNA* acts as a regulatory node, with its suppression promoting NE differentiation, similar to patterns seen in neuroblastoma [48], contributing to genomic instability via increased R-loop formation at transcriptional hotspots. LMNA loss may also explain the heightened replication stress observed in NE SCLC compared to non-NE SCLC, despite its low expression of oncogenes such as *MYC* that typically drive replication stress [3]. Moreover, low *LMNA* expression correlates with poor prognosis in SCLC patients, driven potentially by R-loop-mediated genomic instability [72] or altered differentiation due to R-loop accumulation [73].

Low *LMNA* induced R-loops may be targetable in an array of disease states. For low *LMNA* cancers, specific DNA damaging therapeutics may exacerbate R-loops and associated DNA damage [24], consistent with our data indicating increased DNA damage in siLMNA cells upon topotecan/berzosertib treatment. Conversely, in disease states including laminopathies and aging low *LMNA* induced R-loops could potentially be counterbalanced by agents such as NAT-10 inhibitors which have been shown to ameliorate LMNA mutation induced altered Nup153 recruitment and import/export imbalances [74, 75]. These are critical pathways to explore in future work to determine the potential translational benefit of exploiting the finding of low *LMNA* induced R-loops.

In summary, our study identifies a novel mechanism by which lamin A/C safeguards genome integrity, with important implications for cancer progression and aging. A limitation of this work is the inability to differentiate between the contributions of lamin A and lamin C isoforms. Technical challenges in separately assessing these isoforms currently constrain such analyses, but future studies should explore their specific roles. Additionally, our focus on reduced lamin A/C expression rather than mutations leaves an important avenue unexplored. Investigating LMNA mutations, such as those linked to progeria, could further expand the impact of our findings.

## Materials and Methods

### Cell Lines and Culture Conditions

NCI-H146 (M), NCI-H524 (M), NCI-H69 (M), H69AR (M), NCI-H1048 (F), DMS114 (M) and NCI-H841 (M) cell lines were purchased from ATCC. Cell lines were authenticated using short tandem repeat analysis, and were monthly tested for mycoplasma contamination. Cell medium was RPMI-1640 supplemented with 10% FBS for all lines to maintain consistency. AG04446 and AG12851 (M) fibroblast cells were acquired from participants in the Baltimore Longitudinal Study of Aging from the Coriell Institute Gerentology Research Center and grown in Advanced MEM media supplemented with 15% FBS. Fibroblast lines All four fibroblast cells were immortalized by human Telomerase Reverse Transcriptase protein (hTERT) stable expression using the hTERT Cell Immortalization Kit (ALSTEM, Richmond, CA, USA, cat# CILV02), as previously described [16]. Cell lysates for LMNA protein assessment in SCLC cells were a gift from Dr. Beverly A. Teicher. For EZH2 inhibition experiment NCI-H69 and DMS 273S cells were treated with tazemetostat at 10 μM or control (DMSO) for 27 days with treatment changed every 3-4 days.

### R-loops Immunofluorescence

Briefly, cells were plated at 50,000 cells per well in a 24 well plate on coverslips. Cells were then collected as previously described [77] using the antibody S9.6 (Millipore Sigma #MABE1095). R-loop images were assessed on one of three microscopes Zeiss Axio Observer, Zeiss LSM880 Airyscan, Zeiss LSM780.

### R-loops Slot Blot

Slot blot was performed as previously described [78] using the antibody S9.6 (Millipore Sigma #MABE1095).

### R-loops DRIP-seq

DRIP-seq was performed as previously described [78] using the using the antibody S9.6 (Millipore Sigma #MABE1095).

### Immunoblotting

For immunoblotting the following antibodies were utilized: LMNA/C (Cell Signaling #4777), γH2AX (Millipore Sigma JBW301), EZH2 (Cell Signaling #5246), INSM1 (SantaCruz #sc-271408), GAPDH (Proteintech #60004-1-Ig), α-TUBULIN (Millipore Sigma #T6199), β-ACTIN (Millipore Sigma #A5316), MAb414 (BioLegend #902901), ASCL1 (Cell Signaling #10585).

### Micronuclei Assessment

Cells were plated at 50,000 cells per well in 24 well plates on coverslips. Cells were treated +/-25nM Topotecan for 24 hours followed by a subsequent 24 hours of no treatment. Cells were collected and fixed with 4% PFA for 10 minutes. Cells were stained with DAPI and fixed with Prolong Glass Antifade (ThermoFisher Scientific) and imaged utilizing the Zeiss Axio Observer. For LMNA-KO RNH1 experiments parental and LMNA-KO cells were plated at 250,000 per well in a 6 well plate, the next day they were transfected with either pEGFP-N1-Flag (Addgene #60360) or pEGFP-RNASEH1 (Addgene #108699). The next day they were split into 24 well plates on coverslips, and then allowed another 48 hours to divide prior to collection to assess micronuclei.

### Proximity Ligation Assay

Proximity ligation assay was performed per manufacturer’s instructions utilizing Duolink® reagents (Millipore Sigma #DUO92008, #DUO92004, #DUO92002) and antibodies against S9.6 (Millipore Sigma #MABE1095), and PCNA (SantaCruz #sc-7907). Images were taken using the Zeiss LSM880 Airyscan and analyzed using the Thunderstorm Fiji plugin [79].

### Comet Assay

DMS114 cells were seeded at 250,000 cells per well in a 6 well plate and the subsequent day transfected with siLMNA (Dharmacon #D-004978-03-0010) or siControl (Dharmacon #D-001206-13-20), after 48hours cells were transfected again with either pEGFP-N1-Flag (Addgene #60360) or pEGFP-RNASEH1 (Addgene #108699). Cells were collected after a further 24 hours and processed for the comet assay utilizing the CometAssay Single Cell Gel Electrophoresis Assay (RandD Systems # 4250-050-K) per manufacturer’s instructions. Cells were imaged utilizing the Zeiss Axio Observer, Zeiss LSM880 Airyscan and images were analyzed using the Comet Assay plugin for Fiji based on the NIH Image Comet Assay by Herbert M. Miller from 1997.

### Drug Sensitivity

DMS114 cells were plated at 250,000 cells per well in a 6 well plate and the next day transfected with siLMNA (Dharmacon #D-004978-03-0010) or siControl (Dharmacon #D-001206-13-20) using lipofectamine 3000. After 48 hours cells were split at 2,000 cells per well in a 96 well plate. The following day cells were treated with indicated concentrations of drugs (topotecan and berzosertib), while other cells were frozen for day 0 comparison. After 72 hours the drug treated cells and day 0 were collected and assessed using CellTiter-Glo® (Promgea #G7570) and the EnVision 2104 Multilabel Reader (PerkinElmer).

### Flow Cytometry

DMS114 cells were seeded at 250,000 cells per well in a 6 well plate and the subsequent day transfected with siLMNA (Dharmacon #D-004978-03-0010) or siControl (Dharmacon #D-001206-13-20), after 48hours cells were transfected again with either pEGFP-N1-Flag (Addgene #60360) or pEGFP-RNASEH1 (Addgene #108699). After a further 24 hours ethynyl deoxyuridine (EDU) at 100 μM was added for the last hour prior to collection. EdU was detected using flow cytometry (Click-iT™ EdU Alexa Fluor™ 647 Flow Cytometry Assay, Invitrogen), DNA using DAPI. Data were acquired using a BD LSRFortessa Flow Cytometer and analyzed using FlowJo.

### Plasmids Utilized

The following plasmids were utilized for this project, pEGFP-N1-FLAG (Plasmid #60360), pEGFP-RNASEH1 (Plasmid #108699), pBABE-puro-GFP-wt-lamin A (Plasmid #17662). We also designed a LMNA-KO plasmid, a guide RNA targeting protein coding exons was designed using sgRNA Scorer 2.0 (PMID: 28146356), “GCGCCGTCATGAGACCCGAC**TGG**”, this sequence was annealed and ligated into Lenti-CRISPR-V2 (Addgene# 52961, PMID: 25075903). lentiCRISPR v2 was a gift from Feng Zhang (Addgene plasmid # 52961). This construct was then Sanger sequenced to verify successful guide RNA cloning. This construct has been deposited in Addgene.

### Hematoxylin and Eosin stain (H&E)

DMS 114 Parental vs LMNA-KO cells were plated at 50,000 cells per well in a Lab-Tek II Chamber Slide System (154917). After 24 hours cells were fixed with 4% PFA for 15 minutes, washed in PBS three times, then with 1) Xylene – 10min [3x], 2) 100% EtOH – 3min [2x], 3) 90% EtOH – 3min, 4) 80% EtOH – 3min, 5) 70% EtOH – 3min, 6) Tap Water – 5min, 7) Harris Hematoxylin – 5min, 8) Wash in running tap water – 10min, 9) 3min Eosin, 10) Tap water just to drain excess, 11) 70% EtOH – 10sec, 12) 80% EtOH – 10sec, 13) 90% EtOH – 10sec, 14) 100% EtOH – 10sec, 15) 100% EtOH – 5min, 16) Xylene – 5min [3x], 17) Mount with mounting media (permount). Images were taken using the Keyence BZ-X710.

### EU RNA localization

DMS 114 cells were plated at 250,000 cells per well and then transfected with siLMNA (Dharmacon #D-004978-03-0010), siNXT1 (Dharmacon #L-107194-00-0005), siTPR (Dharmacon #L-010548-00-0005), or siControl (Dharmacon #D-001206-13-20). After 24 hours cells were split at 50,000 cells per well in 24 well plates on coverslips, for parental vs KO and WGA treatment parental and KO cells were split similairly with no previous treatment. EU was added at 2mM for 1 hour and cells were collected after 24 hours and processed using Click-iT™ RNA Alexa Fluor™ 488 Imaging Kit, Invitrogen. For WGA treatment experiments cells were treated +/-2μg/ml WGA and collected after 6 hours. Images were taken utilizing the Zeiss LSM880 Airyscan and quantified utilizing ImageJ. In brief nuclei were delineated utilizing DAPI staining and cytoplasmic border were determined utilizing RNA or tubulin staining. Nuclear signal was subtracted from total signal to determine cytoplasmic RNA signal, with the ratio of the two being utilized to assess RNA localization.

### ACTB-FISH

For ACTB-FISH we utilized the ViewRNA™ Cell Plus Assay Kit (ThermoFIsher Scientific **#**88-19000-99) per manufacturers instructions with DMS 114 Parental and LMNA-KO cells. Images were taken utilizing a Eclipse Ti2 inverted microscope (Nikon). Nuclei were delineated utilizing DAPI staining and cytoplasmic border were determined utilizing Mab414 (BioLegend #902901) staining. Nuclear signal was subtracted from total signal to determine cytoplasmic RNA signal, with the ratio of the two being utilized to assess RNA localization.

### SIM Microscopy

In brief, 250,000 cells per well in a 6 well plate were plated on top of Zeiss High-Performance 0.170mm 18×18 coverslips. Cells were fixed with with 4% Paraformaldehyde in PBS for 15 minutes at RT, permeabilized with 0.5% triton x-100 in PBS, quenched with 0.1M Glycine in PBS for 10 min, washed 3x with TBST (TBS with 0.1% Tween-20) for 10 minutes, blocked in 2% BSA and 0.1% tween 20 in TBS for 60 min, 1st antibody in Blocking Buffer (TBS-T buffer 2% BSA) for 2 hrs at 37C, 1:200 dilution of MAb414 (BioLegend #902901) and 200 ul per sample, wash 3x with TBS-T for 5 minutes, incubated with anti-mouse 488 secondary at 1/400 in blocking solution for 60 min, washed 3x with TBST, mount using Vectashield, and sealed with nail polish. At the same time a separate coverslip with Tetraspeck (ThermoFisher Scientific #T7279) was mounted for SIM standardization.

### Expansion Microscopy

Cells were processed for expansion microscopy according to the protocol described in [80] with following minor change: cells were grown on 12mm cover-glass 1.5H thickness and were placed into glass bottom ibidi dish (Ibidi Cat.No:81158) prior to gelation. Each glass was covered with 1ml of the gelling solution, and then covered with 27mm diameter glass window (Edmun Optics #02-199). After PBS-wash of the denaturation buffer, gels were cut into wedges which fit the lower part of 1.5ml Eppendorf tube vertically. Cells were blocked and stained in a buffer containing 2% BSA, 2% Donkey serum, 0.05% Tween-20, and 0.02% sodium azide in PBS. Anti Nup98 primary antibody (Abcam #ab50610) was used at 1:1000 dilution, for final concentration of 2.4ug/ml. Secondary AffiniPure™ Donkey Anti-Rat IgG (H+L) antibody (Jackson Immune Research #712-005-153) was conjugated in house with Atto-643 dye NHS ester (Atto-tec #AD 643-31) according to suggested protocol https://www.atto-tec.com/fileadmin/user_upload/Katalog_Flyer_Support/Procedures.pdf and used at concentration of 5ug/ml. Concurrent with secondary antibody wash, gels were incubated a few hours with CF568-NHS ester for pan-protein staining and then in YoPro1 (Biotium #40089) at 1uM for nucleic acid staining. For this step, an aliquot of reactive CF568 dye (Biotium #92131, prepared in anhydrous DMSO at the concentration of 5mM dissolved in PBS pH7.4 to the final concertation of 5uM shortly prior to staining.

Water expanded gels were mounted in glass bottom multi-well plates (Cellvis P06-1.5H-N) treated with poly-lysine (Millipore Sigma P4707). To prevent gel drift during imaging, water was added between wells, and the plate was wrapped in parafilm to preserve humidity, and samples were equilibrated on the microscope stage prior to imaging. For better attachment to the glass, gels were gently dried by touching with folded paper – either Kimwipes or lens cleaning paper.

### Multiplexed Immunofluorescence

Multiplex immunofluorescence (mIF) was performed in collaboration with the Human Immune Monitoring Shared Resource (HIMSR) at the University of Colorado School of Medicine. We conducted six-color multispectral imaging using the Akoya Biosciences Vectra Polaris instrument. Unstained formalin-fixed paraffin-embedded (FFPE) slides derived from SCLC tissue were processed on the Leica Bond RX autostainer according to standard protocols provided by Leica and Akoya Biosciences, and routinely performed by HIMSR. Slides from spatially profiled human tumors were consecutively stained with DAPI counterstain and primary antibodies specific to proteins of interest. The primary antibodies — ASCL1, γH2AX, MYC, LMNA, PHH3, and S9.6 — were followed by staining with an HRP-conjugated secondary antibody polymer. HRP-reactive OPAL fluorescent reagents (Opal 570, Opal 690, Opal 480, Opal 620, Opal 780, and Opal 520, respectively) were then applied to deposit fluorescent dyes on the tissue surrounding each HRP molecule. To prevent further dye deposition during subsequent staining steps, slides were stripped between each stain using heat treatment for 20 minutes in Leica Biosystems BOND epitope retrieval solution. Whole-slide scans were collected at 20X magnification. The six-color images were analyzed using inForm software to unmix adjacent fluorochromes, subtract autofluorescence, segment tumor cellular membrane, cytoplasm, and nuclear regions, and score the expression of each protein marker within the relevant cellular compartment. Only the ASCL1, LMNA and S9.6 antibody results were utilized for this project.

### Bioinformatics Analysis

RNA-seq fastq files of parental and LAMN-KO were aligned with hg38 human reference genome using STAR aligner (v 2.7.10b) ([81]). The “–quantMode GeneCounts” module of STAR used to generate raw read counts per gene for RNA-seq dataset. After that, the raw read count matrix was normalized to FPKM. To compare parental and LMNA-KO expression profiles we have performed differential expression analysis using t-tests for RNA-seq dataset. Moreover, ATAC-seq and DRIP-seq/R-loop fastq files were aligned with hg38 reference genome using BWA (v 0.7.17) aligner. The duplicate reads were detected (Picard v3.2.0) and removed before alignment. SAMtools (v 1.19) software was used to index and sort the generated BAM files. After that, MAC2 (v 2.2.7.1) software was utilized for peak calling and BAMScale ‘scale’ module used for generating scaled bigwig files. We used HOMER software “annotatePeaks.pl” PERL script to annotate the genomic regions corresponding to peaks. “multiBigwigSummary”, “computeMatrix”, and “plotHeatmap” modules of deepTools (v 3.5.5) was used to calculate average signal intensity at specific genomic regions and to generate heatmaps. ATAC seq peaks were assigned by priority to: 1) Promoter if they were within 2kb of a TSS 2) Intragenic if peaks overlapped with a gene body or 3) intergenic if they were not assigned to one of these categories. The nearest approach was utilized i.e. peaks were only assigned to one category and one gene, with most proximal TSS taking precedence. Peaks were then assessed for increased accessibility in either Parental or LMNA-KO DMS114 cells using a threshold of p<0.05 (Right). Gene set enrichment analysis was performed using ssGSEA/GSEA with “msigdb” and “GSVA” libraries in R (v 4.2.3). IGV (Integrative Genomics Viewer) tool (version 2.16.0) used for the visualization. “matplotlib.pyplot” python (version 3.9.17) library used to generate volcano plots, and lollipop plots.

### Patient Survival Analysis

Gene expression data for the chemo-naive SCLC cohort ([59]) was obtained from cBioPortal ([82]), which included clinical data. Single-gene survival analysis was performed by separating patients into high and low groups based on expression, followed by survival analysis using the survival (v3.6-4) and survminer (v0.4.9) packages in R (v4.2.3). In addition, single sample gene set enrichment analysis was also calculated for chromosome attachment genes and R-loop genes using the corto (v1.2.4) R package [83]. These values were used to separate patients with high and low gene set activity, followed by survival analysis.

