## Supplemental Figures and Tables for "Lamin A/C Deficiency Drives Genomic Instability and Poor Survival in Small-Cell Lung Cancer through Increased R-loop Accumulation"

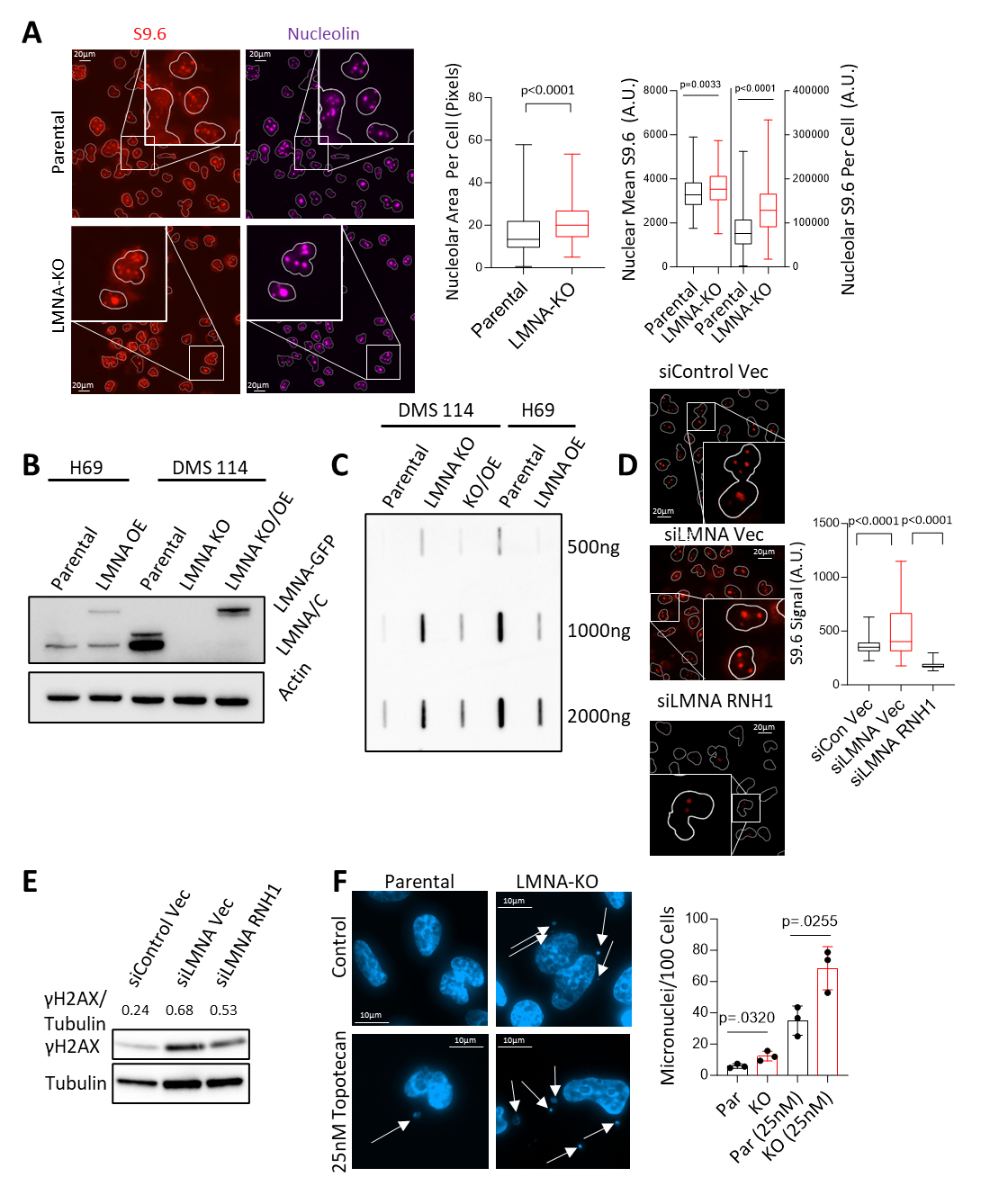

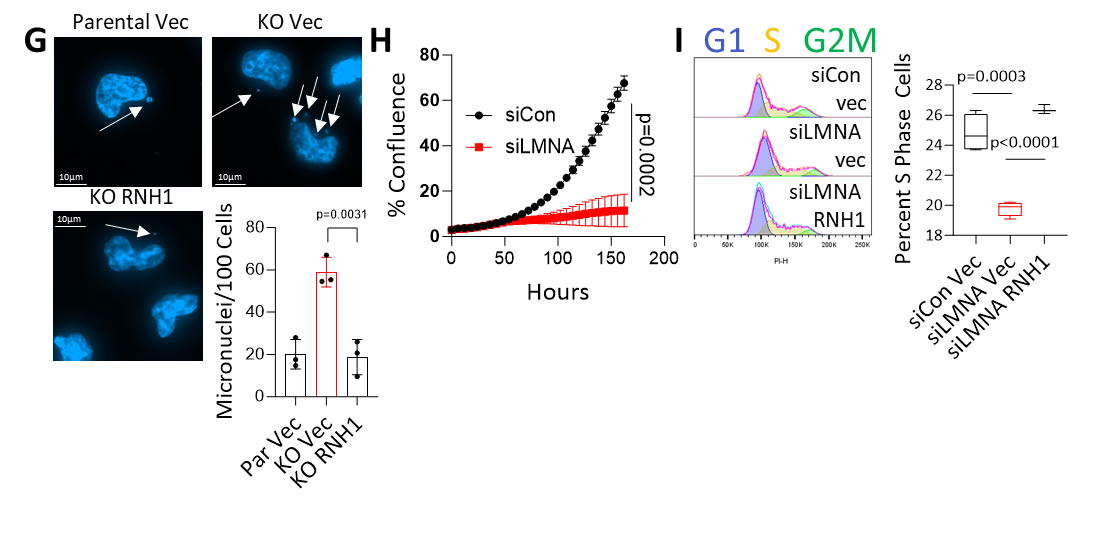

**Figure S1:** **A)** Immunofluorescence of R-loops and nucleoli in parental and *LMNA*-KO DMS114 SCLC cells using S9.6 and nucleolin antibodies. **B)** Expression of *LMNA* and the LMNA-GFP construct in NCI-H69 and DMS114 cells. **C)** Slot blot of R-loops in *LMNA*-KO and LMNA-GFP overexpressing (OE) NCI-H69 and DMS114 cells compared to their parental counterparts. **D)** Immunofluorescence assessment of R-loops stained with S9.6 in siLMNA and siControl DMS114 cells, with RNaseH1-overexpression (OE) control. **E)** Immunoblotting of γH2AX to evaluate DNA damage in siLMNA and siControl DMS114 cells, with RNaseH1-OE control. **F,G)** Representative images and quantification of micronuclei in DMS114 parental and LMNA-KO cells, (F) with or without treatment with 25nM of topotecan, (G) or with RNaseH1-OE control. White arrows indicate micronuclei. **H)** Growth rate of siLMNA and siControl DMS114 cells assessed using Incucyte. **I)** Percentage of DMS114 cells in S phase as determined by flow cytometry of DNA content in siLMNA and siControl cells with RNaseH1-OE control.

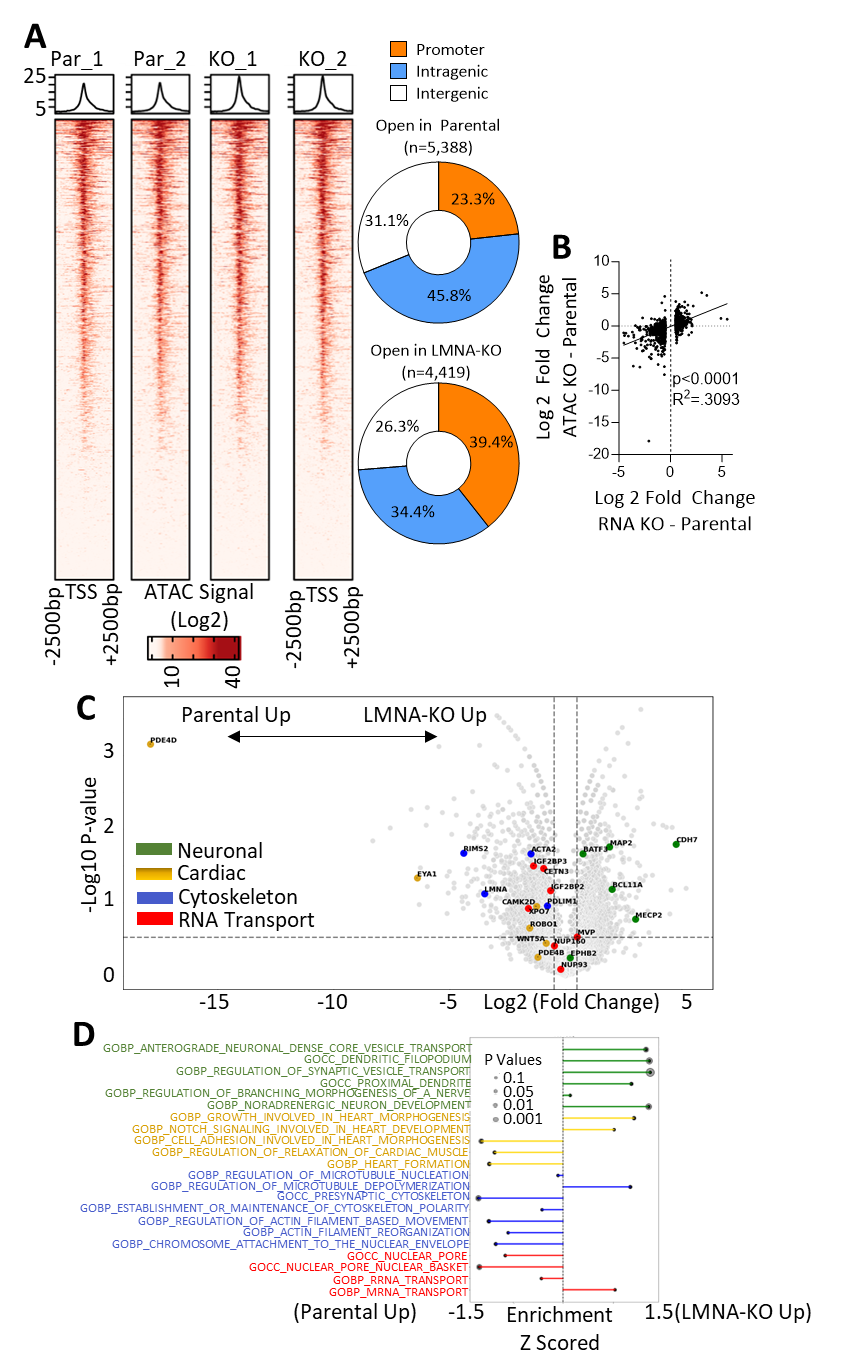

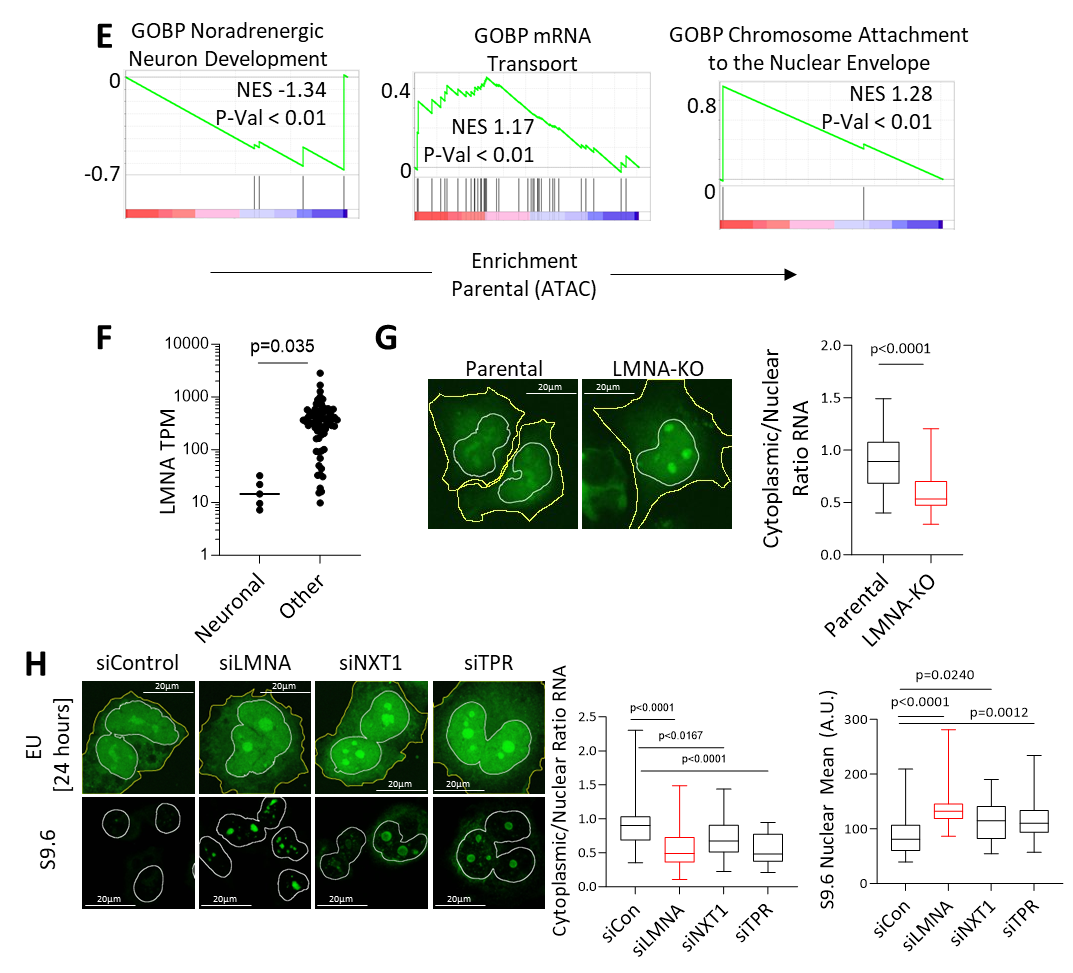

**Figure S2**: **A)** Normalized ATAC-seq signal intensity proximal to the transcription start site (TSS) +/- 2500bp (left) and percentage of peaks based on genomic location i.e. promoter, intragenic, intergenic (right). **B)** Correlation between ATAC promoter accessibility and RNA expression in genes with ≥ 0.5 log2 fold change differential expression between parental and *LMNA*-KO DMS114 cells. **C)** Volcano plot showing promoter accessibility by gene in *LMNA*-KO vs parental DMS114 cells (ATAC-seq). **D)** Enrichment of pathways in *LMNA*-KO vs parental DMS114 cells as determined by ssGSEA of ATAC-seq promoter accessibility. **E)** Leading edge genes from Fig. 2C assessed utilizing ATAC-seq promoter accessibility by GSEA. **F)** Single cell RNA-seq data from the Human Protein Atlas showing average *LMNA* RNA expression (TPM) from 689,601 cells of non-diseased human tissues, broken into neuronal lineages (inhibitory, excitatory and bipolar neurons, rod and cone photo receptors) vs. all other lineages. **G)** RNA export capacity assessed utilizing EU incorporation in parental and *LMNA* KO DMS114 cells, nuclei are outline in white while the cytoplasm is outlined in yellow. **H)** RNA export and R-loop formation assessed following silencing of NXT1, TPR, and lamin A/C. RNA export was measured by EU incorporation, and R-loops detected by immunofluorescence, representative images are shown with nuclei outlined in white while the cytoplasm is outlined in yellow.

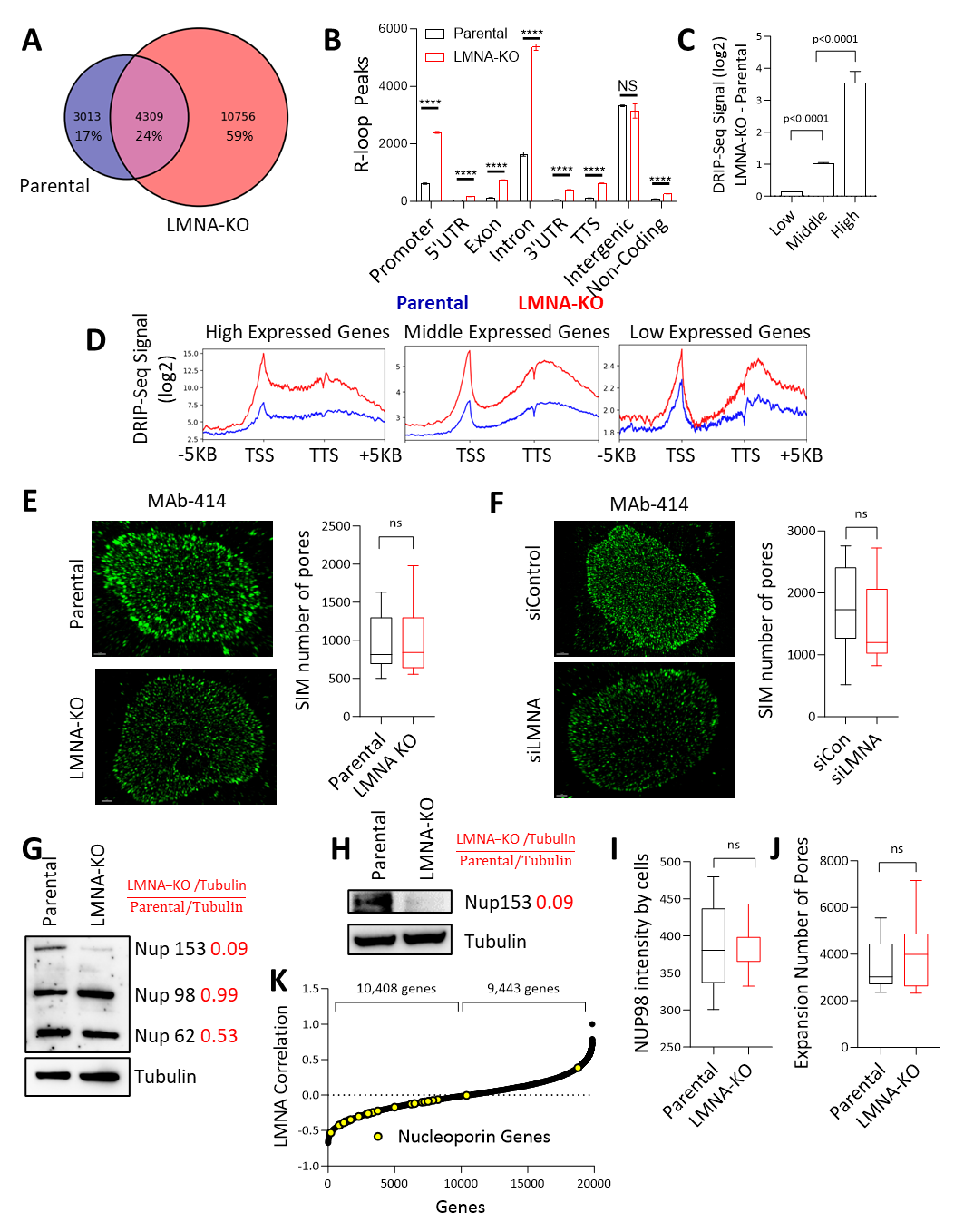

**Figure S3: A)** DRIP-seq signal peaks in parental or *LMNA*-KO DMS114 cells. **B)** Analysis of R-loop peaks across parental and *LMNA*-KO DMS114 cells, with peaks categorized by the most proximal transcription start site (TSS). Numbers indicate fold changes in R-loop peaks between *LMNA*-KO and parental settings. **C)** DRIP-seq signal mapped to the genome, and total R-loop signal across each gene assessed in a strand-specific manner. Genes were categorized as up-regulated in either parental or *LMNA*-KO DMS114 cells, using thresholds of p<0.05 and an absolute ≥2-fold change in DRIP-seq signal. Genes not meeting these criteria were designated as unaltered. **D)** R-loop levels per gene determined utilizing DRIP-seq, and genes were split into, high (>6 log2 TMM FPKM, 447 genes), middle (1-5 log2 TMM FPKM, 10,045 genes) and low (<1 log2 TMM FPKM, 27,290 genes) expression categories based on overall expression based on RNA-seq data. DRIP-seq signal differences were averaged by category and compared between parental and *LMNA*-KO settings across gene bodies. **E,F)** MAb414 staining of individual nuclear pores visualized using SIM in *LMNA*-KO (E) and siLMNA (F) cells versus controls using DMS114 cells. **G)** FG nucleoporins assessed by immunoblotting using MAb414 staining in parental and *LMNA*-KO DMS114 cells. **H)** Nup-153 assessed by immunoblotting in parental and *LMNA*-KO cells. **I,J)** Expansion microscopy with NUP98 immunofluorescence used to assess NPCs, specifically examining (I) NUP98 staining intensity and (J) the number of NPCs per cell. **K)** LMNA RNA expression across the CCLE panel of 1040 cell lines vs all other genes.

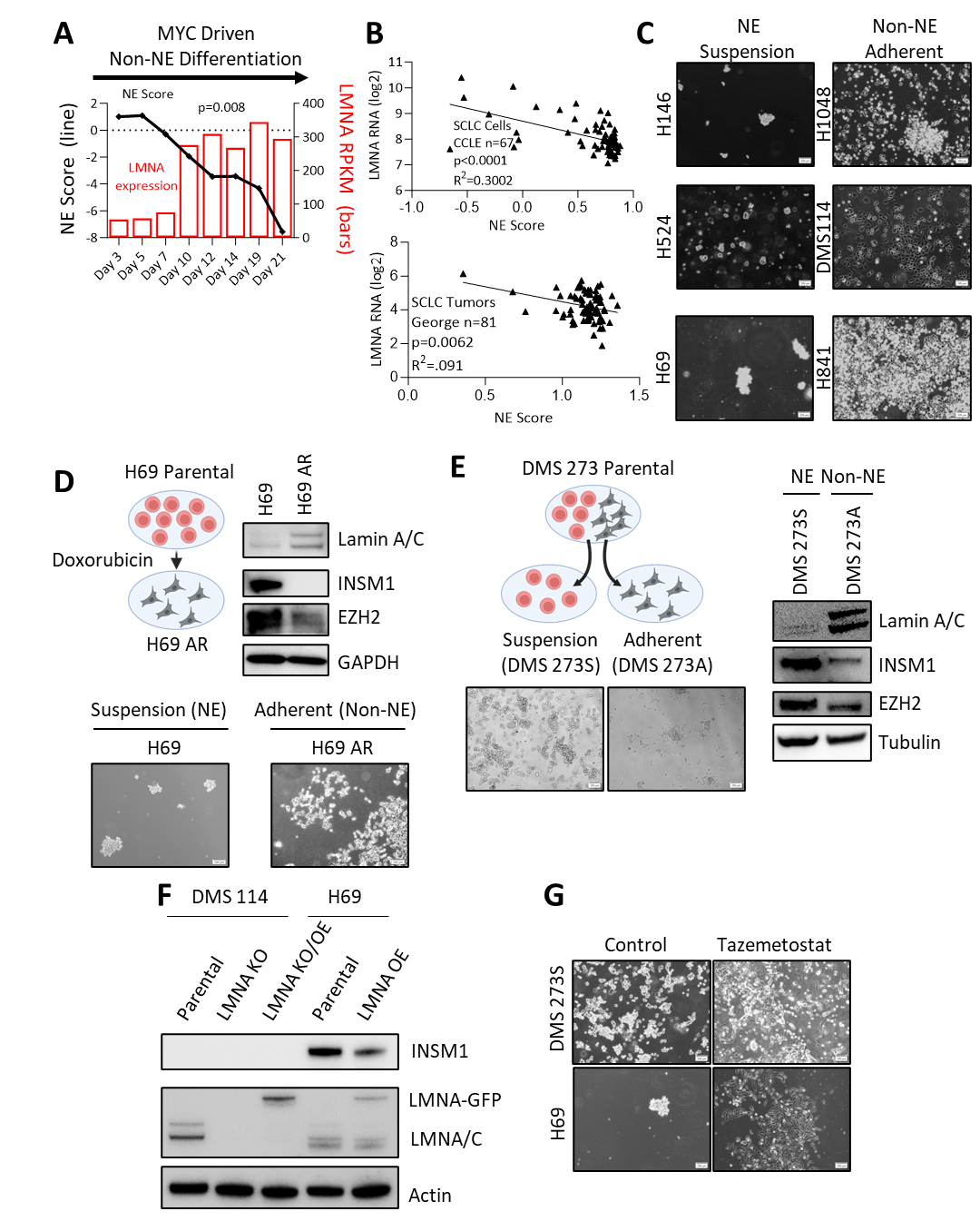

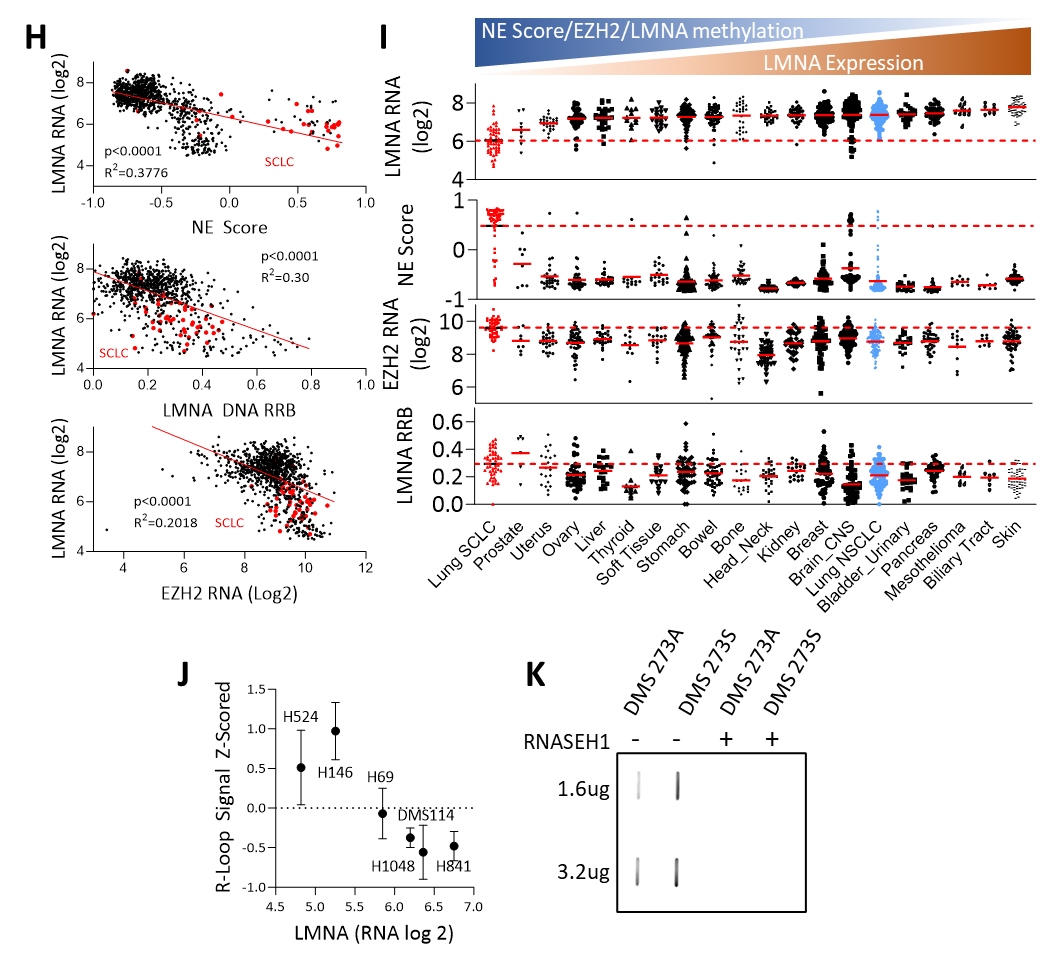

**Figure S4**: **A)** NE differentiation and LMNA expression over time during NE to non-NE transition in a MYC and NOTCH-driven model [51] **B)** Correlation between LMNA RNA expression and NE differentiation across SCLC cell lines (CCLE) and SCLC patient tumors [59]. **C)** Brightfield images of SCLC cell lines. **D)** Brightfield images of alongside western blot assessment of INSM1, lamin A/C, and EZH2 in parental NCI-H69 (suspension, NE), and non-NE derivative H69AR (doxorubicin-treated, adherent). **E)** Brightfield images alongside western blot assessment of INSM1, lamin A/C, and EZH2 in of DMS273S (suspension) and DMS273A (adherent) cell lines. **F)** Immunoblot analysis of lamin A/C and INSM1 in DMS114 and NCI-H69 isogenic LMNA models. **G)** Brightfield images of DMS273S and NCI-H69 cells treated with EZH2 inhibitor tazemetostat. **H)** LMNA expression (microarray) plotted against, NE differentiation, EZH2 expression (microarray), and LMNA gene promoter methylation (DNA RRBS methylation) across the CCLE, SCLC cell lines are marked in red. **I)** LMNA expression (microarray), NE differentiation, EZH2 expression (microarray), and LMNA gene promoter methylation (DNA RRBS methylation) by cancer type across the CCLE, with a dotted red line representing average expression in SCLC cells. **J)** R-loop assessment from representative blot (Fig. 4D) against *LMNA* RNA expression. **K)** R-loop assessment in DMS273 adherent and suspension cell lines using slot-blot, with RNase H1 treatment as a control.

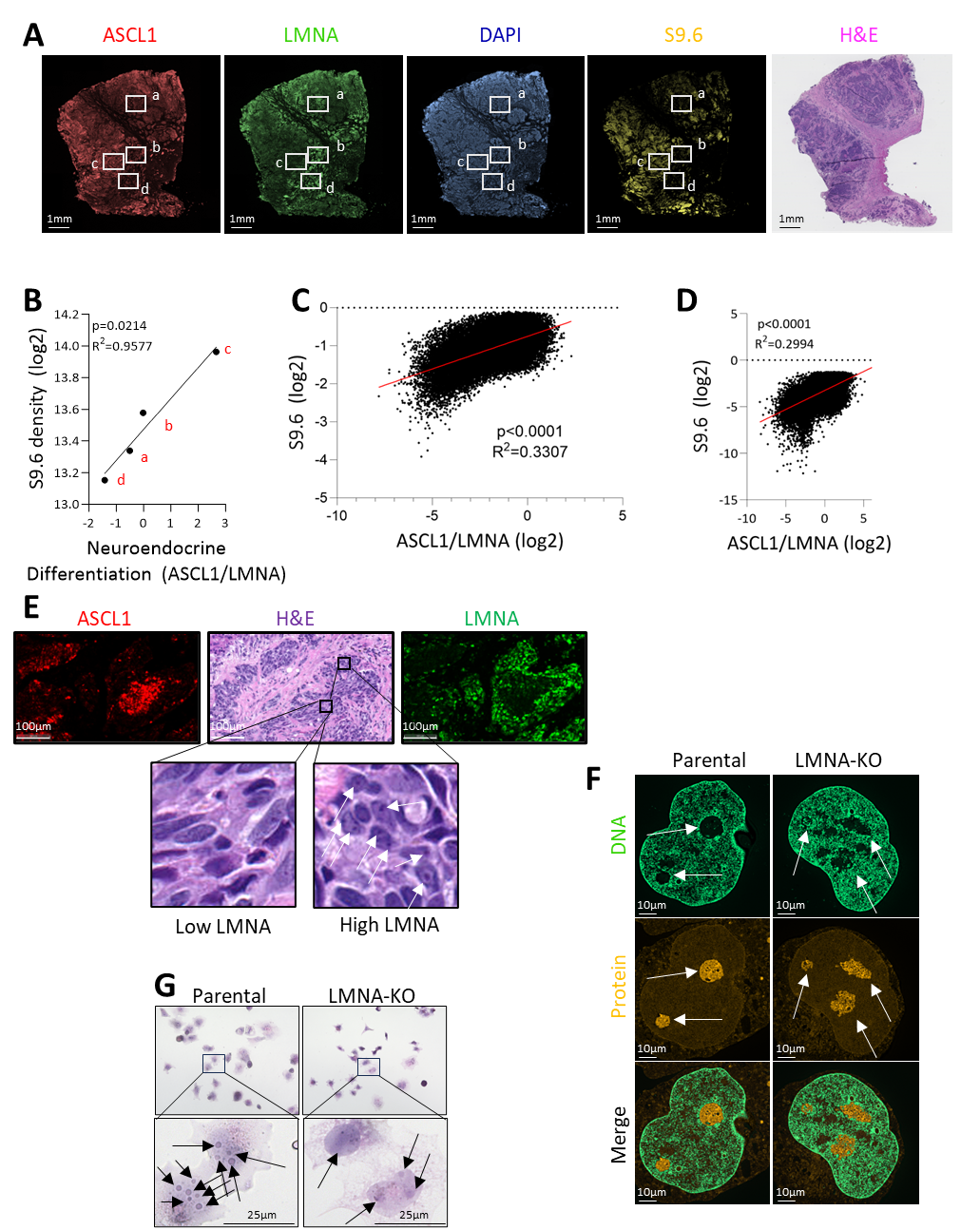

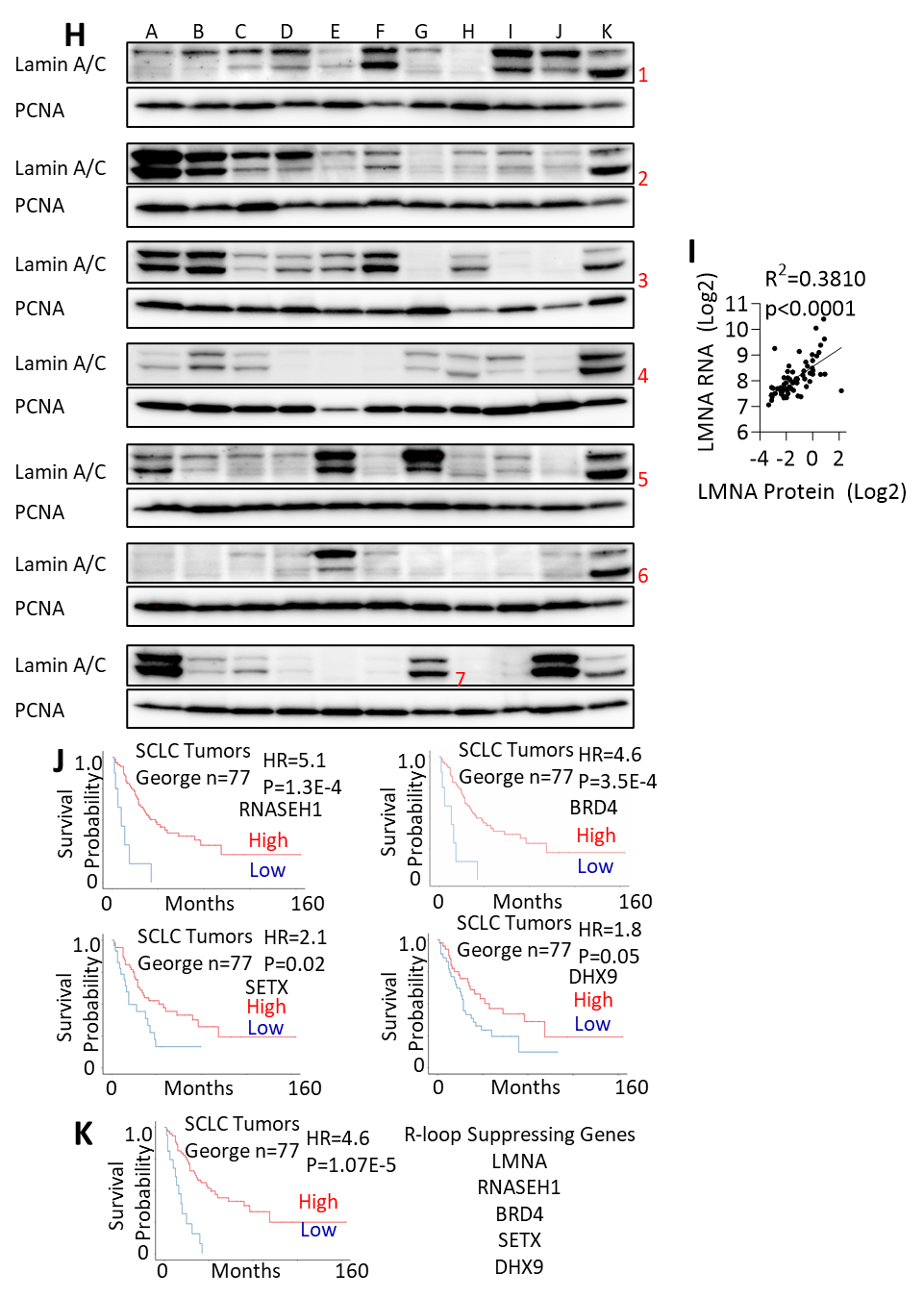

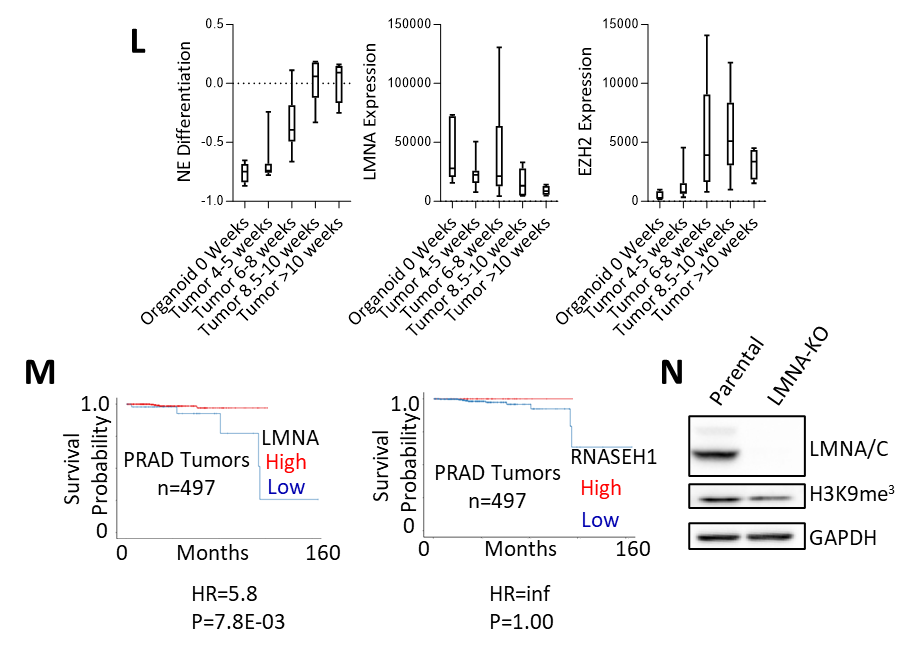

**Figure S5: A)** SCLC rapid autopsy sample (RA-17) stained for DAPI, ASCL1, LMNA, R-loops, and H&E. Four representative regions of interest are shown in white boxes marked a-d. **B)** Relationship between ASCL1/LMNA signal and R-loops from regions of interest from panel A, marked a-d. **C)** Relationship between ASCL1/LMNA signal and R-loops from single cells across all four regions of S5A. **D)** Relationship between ASCL1/LMNA signal and R-loops from the TMA regions (representative images 5C). **E)** Representative H&E staining of RA-17. Arrows indicate nucleoli. **F)** The perinucleolar boarder, nucleolar clearing and nucleolar protein density assessed using expansion microscopy in parental and LMNA-KO DMS114 cells. Arrows indicate example nucleoli. **G)** DMS114 parental and LMNA-KO cells stained with H&E. Arrows indicate example nucleoli. **H)** Lamin A/C protein expression assessed across 63 SCLC cell lines by western blot utilizing PCNA as a control per each blot and DMS 114 as a control to maintain consistency between blots. **I)** Correlation between LMNA protein expression (from S5H) and RNA expression (available from CellMiner CDB). **J)** Expression of R-loop suppressing agents and overall survival in SCLC patients [59]. **K)** ssGSEA analysis of a panel of R-loop suppressing genes (LMNA, RNASEH1, BRD4, SETX, DHX9) and overall survival SCLC patients [59]. **L)** NE differentiation status (assessed by ssGSEA) alongside LMNA and EZH2 RNA expression over various timepoints from 10 patient prostate cells transformed using the PARCB model [62]. **M)** Expression of LMNA and R-loop suppressing agent RNASEH1 and overall survival from TCGA PRAD patients data. **N)** H3K9me3 and lamin A/C assessed by immunoblot in in parental and LMNA-KO DMS114 cells.

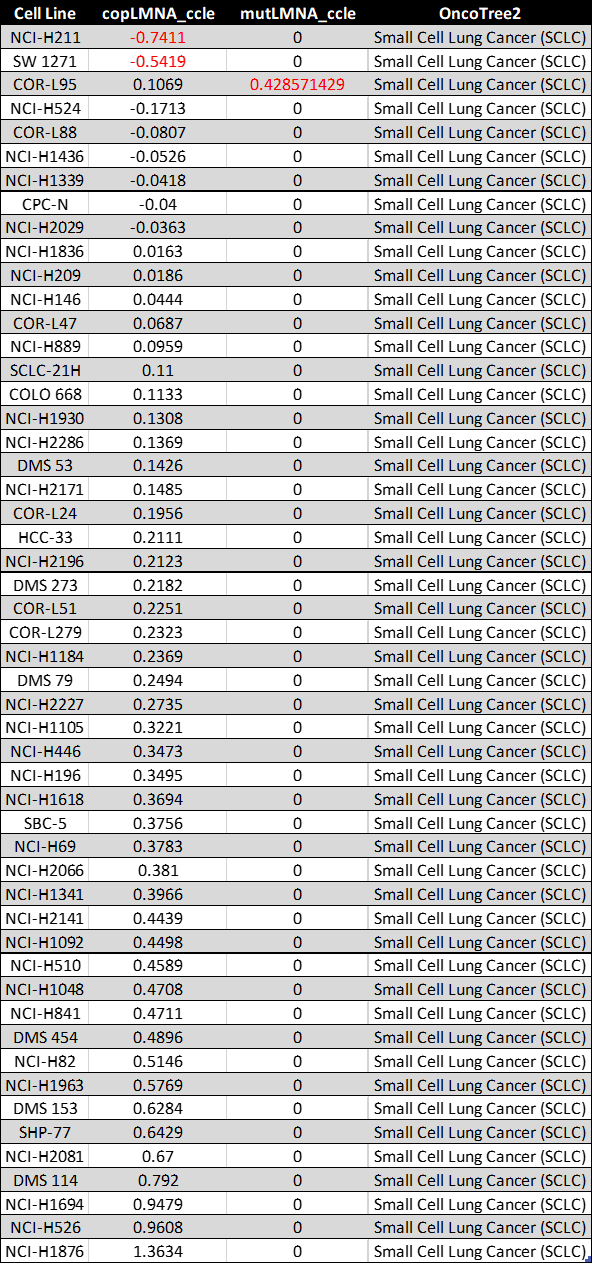

**Supplemental Table 1:** LMNA Mutation and copy number data for SCLC cells lines from the CCLE dataset. Cell lines with note-worthy copy number loss and/or mutation of LMNA are highlighted in red. These data are publicly available using CellMiner [76].

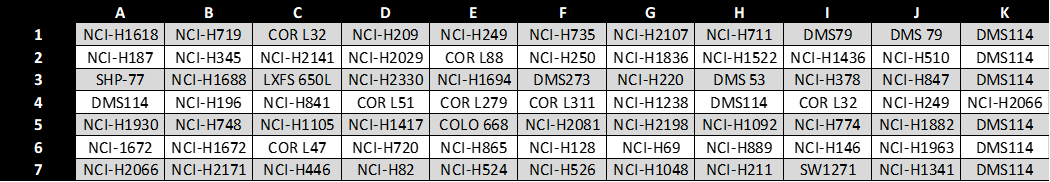

**Supplemental Table 2:** This key is in reference to Supplemental Figure 5E wherein lamin A/C protein expression was assessed across 63 SCLC cell lines by western blot utilizing PCNA as a control per each blot and DMS114 as a control to maintain consistency between blots.
